# Plasticity in nonsense-mediated decay and translation initiation regulate polyphenism

**DOI:** 10.64898/2026.03.27.714762

**Authors:** Penghieng Theam, Hanh Witte, Rebecca Liu, Tobias Loschko, Christian Rödelsperger, Cátia Igreja, Ralf J. Sommer

## Abstract

Developmental plasticity is increasingly recognised as facilitator of evolutionary novelty. However, how plasticity itself evolves and how variation in plastic trait expression is structured in populations remain unknown^1,2^. The predatory nematode *Pristionchus pacificus* exhibits mouth-form plasticity with underlying molecular mechanisms being increasingly identified^3^. We investigate the temporal scale of natural variation of mouth-form plasticity. An 11-year survey characterised *Adoretus* beetle-derived isolates from Colorado, La Réunion Island and revealed a gradual shift in mouth-form preference. Quantitative trait locus mapping of mouth-form preferences identified a single peak harbouring the developmental switch gene *eud-1*. Through CRISPR-engineering and biochemical assays, we show that plasticity in nonsense-mediated decay coupled with alternative start codon selection resulting in different N-terminal proteoforms of EUD-1 are associated with natural variation of mouth-form preference. This work provides molecular explanations for variation in plastic trait expression and links nonsense variants in the major developmental switch locus to ecological and evolutionary processes.

## Background

Resource polyphenisms are some of the most extreme examples of developmental plasticity and are known to facilitate the utilisation of distinct food sources in many animals^1^. In general, developmental (phenotypic) plasticity, the ability of a genotype to produce different phenotypes in response to environmental variation, is increasingly recognised as a major facilitator of evolutionary novelty and organismal diversification^2^. Recent case studies are beginning to identify gene regulatory networks (GRNs) in vertebrates, insects, nematodes and other organisms exhibiting polyphenisms and other plastic traits^3^. Such molecular understanding on how plastic systems work is crucial for an overall acceptance of developmental plasticity as a significant factor for evolution. However, scepticism remains and important research questions about the relationship between developmental plasticity and evolution are still unanswered. For example, little is known about how plasticity itself evolves and how variation in plastic trait expression is structured in natural populations.

One animal that has been intensively studied in the last decade is the facultative predatory nematode *Pristionchus pacificus,* which exhibits two alternative mouth forms resulting in different feeding strategies^4,5^. During postembryonic development, individual *P. pacificus* worms execute one of two alternative morphs in an irreversible manner^6^. Stenostomatous (St) animals have a single tooth and a narrow mouth (called the buccal cavity in nematodes) and are strict bacterial feeders (Fig. 1a). In contrast, eurystomatous (Eu) animals have two teeth with a broad buccal cavity and can supplement their bacterial diet by predation on other nematodes (Fig. 1a). This mouth-form dimorphism is a bistable developmental switch with various environmental stimuli being able to influence the mouth-form ratio, making it an attractive system to manipulate plastic trait formation^7–12^. The GRN controlling mouth-form development has been studied in detail and the sulfatase-encoding gene *eud-1* has been identified as the key developmental switch locus^13–19^ (Fig. 1b). EUD-1 acts as a dosage-dependent switch and mutations in *eud-1* result in the absence of the Eu mouth form^13^. However, the characterisation of a GRN on its own does not provide evidence for the ecological and evolutionary significance of a plastic trait or developmental plasticity in general^20^. Such claims, and further arguments for the potential adaptive value of developmental plasticity require specific ecological and evolutionary investigations^21^. Here, we study the temporal scale of natural variation of mouth-form plasticity and made the surprising finding that plasticity in nonsense-mediated decay (NMD) and translational control of the key developmental switch gene *eud-1* regulate natural variation in mouth-form bias.

**Fig. 1.**
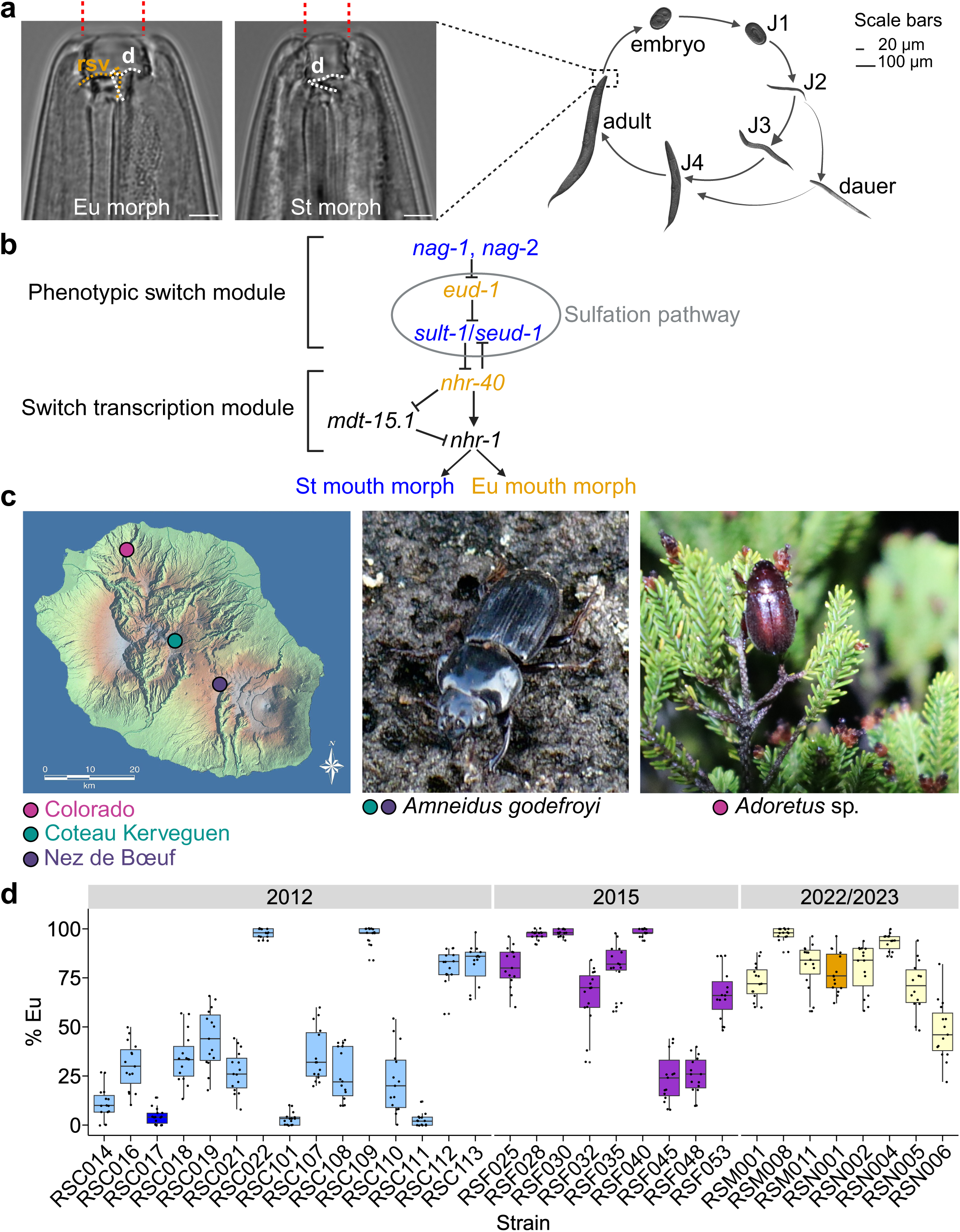
*Adoretus* beetle-derived *P. pacificus* strains provide St-biased material for mouth-form natural variation studies. **a,** Nematode life cycle (right) and Nomarski micrographs of adult head structures (left). Adult hermaphroditic *P. pacificus* individuals have an irreversible feeding polyphenism. They can either possess just one dorsal (d) tooth and feed strictly on bacteria (St morph) or decorate their right subventral (rsv) sector of the buccal cavity with another tooth enabling them to kill other nematodes and become omnivorous feeders (Eu morph). **b,** The mouth-form decision is controlled by a complex gene regulatory network with various gene modules, including a ‘phenotypic switch module’ and a more downstream ‘switch transcription module’. Central to the ‘phenotypic switch module’ is the sulfation pathway comprising the sulfatase-encoding gene *eud-1* and the sulfotransferase-encoding gene *sult-1*/*seud-1*. *eud-1* is the main mouth-form switch gene and *P. pacificus* adults become St when *eud-1* is mutated. Genes shown in blue and orange result in all-Eu and all-St phenotypes, respectively. **c,** Map of La Réunion Island with scarab beetle sample localities indicated. Among other scarab beetles, *P. pacificus* can be isolated from the endemic stag beetle *Amneidus godefroyi* collected from Nez de Bœuf (violet dot) and Coteau Kerveguen (green dot) and species of the genus *Adoretus* collected from Colorado (magenta dot). **d,** Natural variation of mouth-form preference of *Adoretus*-derived *P. pacificus* strains collected from Colorado in different years including 2012 (light blue and blue), 2015 (purple) and 2022 and 2023 (light yellow and orange). Boxplots show the median (centre line), interquartile range (IQR; box), and whiskers extending to 1.5×IQR of mouth-form ratio (*n* = 15 replicates). Closely-related strains RSC017 (St-biased; blue) and RSN001 (Eu-biased; orange) were selected for further studies (Suppl. Fig. 1). Source data are provided as a Source Data file. Scale bars: 20 µm for embryo and J1 stages, 100 µm for the rest of the stages (right), and 5 µm for mouth-form micrographs (left) (**a**) and 20 km for map of La Réunion (**c**). Map of La Réunion. © Conseil Régional de La Réunion (2006) at Wikimedia Commons, CC BY-SA 2.5. *A. godefroyi* photo. © Dr. Matthias Herrmann (**c)**.

### *Adoretus* beetle-derived *P. pacificus* strains are a hotspot for stenostomatous-biased genotypes

The overwhelming majority of *P. pacificus* natural isolates from around the world preferentially express the Eu mouth form under standard laboratory growth conditions^13,22^. We performed large-scale surveys using the Sommer Lab strain collection with more than 2,000 *P. pacificus* natural isolates in search for genotypes that would be biased towards the St morph instead^23–27^. Ideally, we aimed for pairs of *P. pacificus* strains from the same locality and even the same beetle host species with opposite mouth-form preferences. Indeed, we found that in material isolated in 2012 from beetles of the genus *Adoretus* at Colorado on La Réunion Island most strains are St-biased (Fig. 1c,d). Colorado is in the northern part of the Island close to the capital St. Denis at an altitude of around 600 m with mixed, endemic and invasive vegetation (Fig. 1c). Of 15 *P. pacificus* strains isolated in 2012, 11 strains were preferentially St: four strains were below or around 10% Eu, whereas the other strains were 10-40% Eu. The remaining four strains were Eu-biased (Fig. 1d). Thus, the majority of strains isolated from *Adoretus* beetles at Colorado are St-biased under laboratory conditions.

We performed additional surveys at Colorado in 2015 and in 2022 and 2023. From a smaller survey in 2015, seven of nine isolates were Eu-biased and only two strains were preferentially St (Fig. 1d). In two more recent samplings in 2022 and 2023, all eight newly isolated strains were highly Eu (Fig. 1d). Note that 2022 and 2023 were extremely dry years, which resulted in low *P. pacificus* yields although the number of *Adoretus* beetle samples were quite large. Taken together, *Adoretus*-derived *P. pacificus* sampling at one particular locality on La Réunion Island identified a shift in mouth-form preferences from St-biased towards Eu-biased strains over an 11-year period and resulted in the isolation of a total of 13 wild isolates with a mouth-form ratio biased towards the St form, which is otherwise rare throughout the world. Note that we have initiated ongoing studies on an annual basis to better understand the dynamics and fluctuations of *Adoretus*-derived *P. pacificus* strains from Colorado.

### Recombinant inbred line analysis identifies a single QTL at the *eud-1* locus

Genome-wide single nucleotide polymorphism (SNP) analysis indicated that the *Adoretus*-derived strains from Colorado are indeed very closely related to each other (Suppl. Fig. 1). To study the molecular composition associated with the natural variation in mouth-form preference of strains from Colorado, we selected *P. pacificus* RSC017 (a St strain isolated in 2012) and RSN001 (a Eu strain isolated in 2023) to perform a recombinant inbred line (RIL) experiment. We followed the protocol previously established in *P. pacificus*^28–30^ (Fig. 1d and 2a). In short, after reciprocal crosses between RSC017 and RSN001, we inbred 141 lines until the F12 generation and performed genotyping and phenotyping between generations 13-15 (Suppl. Fig. 2) (see Methods for details). Quantitative trait locus (QTL) analysis revealed a single locus on the X chromosome in the region of *eud-1* (Fig. 2b,c). Strikingly, a previous study of *P. pacificus* strains from a different clade and a different locality on Réunion Island had also identified a single QTL at the *eud-1* locus^28^ (Fig. 1c and Suppl. Fig. 1). Thus, RIL analysis of *Adoretus*-derived strains RSC017 and RSN001 identified a single QTL at the *eud-1* locus indicating this gene to be repeatedly associated with natural variation in mouth-form preference.

**Fig. 2.**
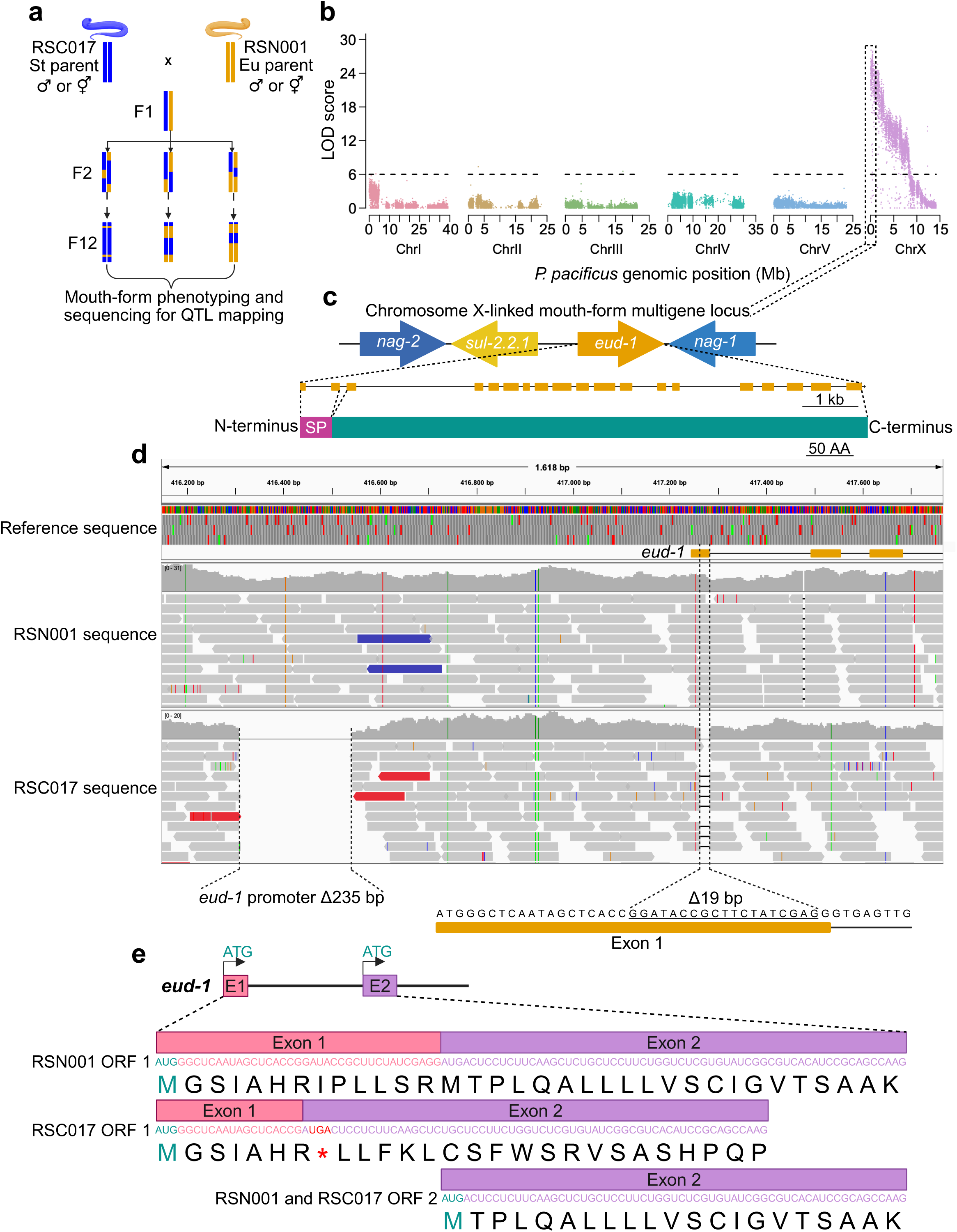
Recombinant inbred line (RIL) analysis revealed a single quantitative trait locus (QTL) at the *eud-1* gene. **a,** Schematic of RIL generation. Reciprocal cross between self-sperm-depleted hermaphrodites of one strain with males of the other strain and vice versa. This panel of RILs was inbred until generation F12 in which near homozygosity is achieved. Mouth-form phenotyping and sequencing were carried out between generations F13-F15. The mouth-form phenotype of RILs is shown in Suppl. Fig. 2. **b,** QTL mapping revealed a single significant peak on chromosome X. A logarithm of the odds (LOD) score above six indicates a significant association. The most significant region spanned ∼0.4 Mb on the left arm of chromosome X and corresponded to the mouth-form supergene locus. **c,** The supergene locus contains four genes, three of which (*nag-1*, *nag-2* and *eud-1*) contribute to the mouth-form decision. *eud-1* is epistatic over the two *nag*-genes. Below the gene organization, an enhanced representation of the exon-intron structure of *eud-1* and a simplified protein structure are shown. The first two exons of *eud-1* encode a signal peptide (SP) important for the proper subcellular localisation of EUD-1 protein (Fig. 6b). **d,** *eud-1* locus 5’ vicinity and exon 1 of RSN001 and RSC017 genomes compared to the reference *P. pacificus* genome (version El Paco) viewed on the software IGV. Genomic position is shown on the top. In the RSN001 and RSC017 genome panels, arrow blocks represent strand-oriented reads aligned to the reference genome. Read coverage profiles are represented by histogram-like tracks above each genome. Polymorphisms between RSN001 and RSC017 genomes and the reference genome including SNPs and deletions are indicated by coloured lines and gaps on the read coverage tracks, respectively. Two deletions in the *eud-1* locus were observed in the St-biased RSC017 strain. One deletion spans 235 bp in the promoter region and is located 701 bp upstream of the coding sequence. The other deletion removes 19 bp in the first exon (underlined). **e,** Representation of the first and second exons of *eud-1* and alternative coding potentials. The 19 bp deletion in exon 1 of the RSC017 strain leads to a frameshift mutation and a premature stop codon (red asterisk). A second in-frame open reading frame (ORF) can start from the second exon in both strains. Scale bars: 1 kb and 50 AA for *eud-1* gene and EUD-1 protein structures, respectively (**c**).

### Sequencing of St-biased strains identified molecular lesions in *eud-1* coding sequences

We performed whole genome sequencing of RSC017 and RSN001 and all other St-biased strains to identify potential molecular lesions in *eud-1* and its flanking genes. The *eud-1* locus was previously shown to form a supergene with two other genes regulating mouth-form plasticity^15^ (Fig. 2c). We found two different variants at the *eud-1* locus in these St-biased wild isolates, both of which are different from those previously identified^28^. First, we found a variant in the promoter region of *eud-1* with the St-biased strain RSC017 having a 235 bp deletion in the promoter, located 701 bp upstream of exon 1 (Fig. 2d). Interestingly, this deletion was found in 18 additional strains and was present in the meta-population between 2012 and 2023 (Supplementary Table 1). Second, RSC017 harbours a nonsense *eud-1* variant with a 19 bp deletion in the coding sequence (CDS) of exon 1, which encodes part of the signal peptide of EUD-1 (Fig. 2c,d). This 19 bp deletion results in a frameshift (Fig. 2d,e). Note that this deletion is close to the boundary of exon 1 and intron 1 and creates a new *cga* codon (Arg) with the final ‘*a*’ nucleotide coming from exon 2, which normally starts with an *aug* (Met) codon (Fig. 2d,e). This 19 bp deletion was found in seven more independent strains isolated between 2012 and 2015 (predominantly St), but it was not found in any strains isolated in 2022 and 2023 (Supplementary Table 1). We also searched for natural variants in the neighbouring genes that have been implicated in mouth-form regulation, but did not find any natural variants that would be considered as candidates. Thus, a novel RIL experiment and whole genome sequencing of the 13 St-biased strains identified two new molecular variants at the *eud-1* locus indicating that mutations in this gene are over-represented in nature.

### The *eud-1* promoter variant is not involved in mouth-form regulation

In *P. pacificus*, CRISPR-associated engineering has been well-established and used in various large-scale studies^31,32^. We used CRISPR engineering to test for the functional significance of natural variants observed between RSC017 and RSN001 (Fig. 3a–c and Supplementary Tables 2 and 3). In principle, RSC017-specific deletions can be eliminated in the RSN001 background, or in a reverse way, RSN001-specific sequences can be reintroduced into the RSC017 background (Fig. 3a–c). First, we investigated a potential role of the 235 bp deletion in the RSC017 *eud-1* promoter region. Specifically, we deleted the 235 bp fragment that is absent in RSC017 in the Eu-biased strain RSN001 using a repair template (Supplementary Table 3). We found that two independent lines with a 235 bp deletion in the RSN001 genetic background had a high Eu mouth-form ratio similar to wild type RSN001 animals (Fig. 3a). We conclude that this natural variant in the *eud-1* promoter does not on its own control mouth-form expression.

**Fig. 3.**
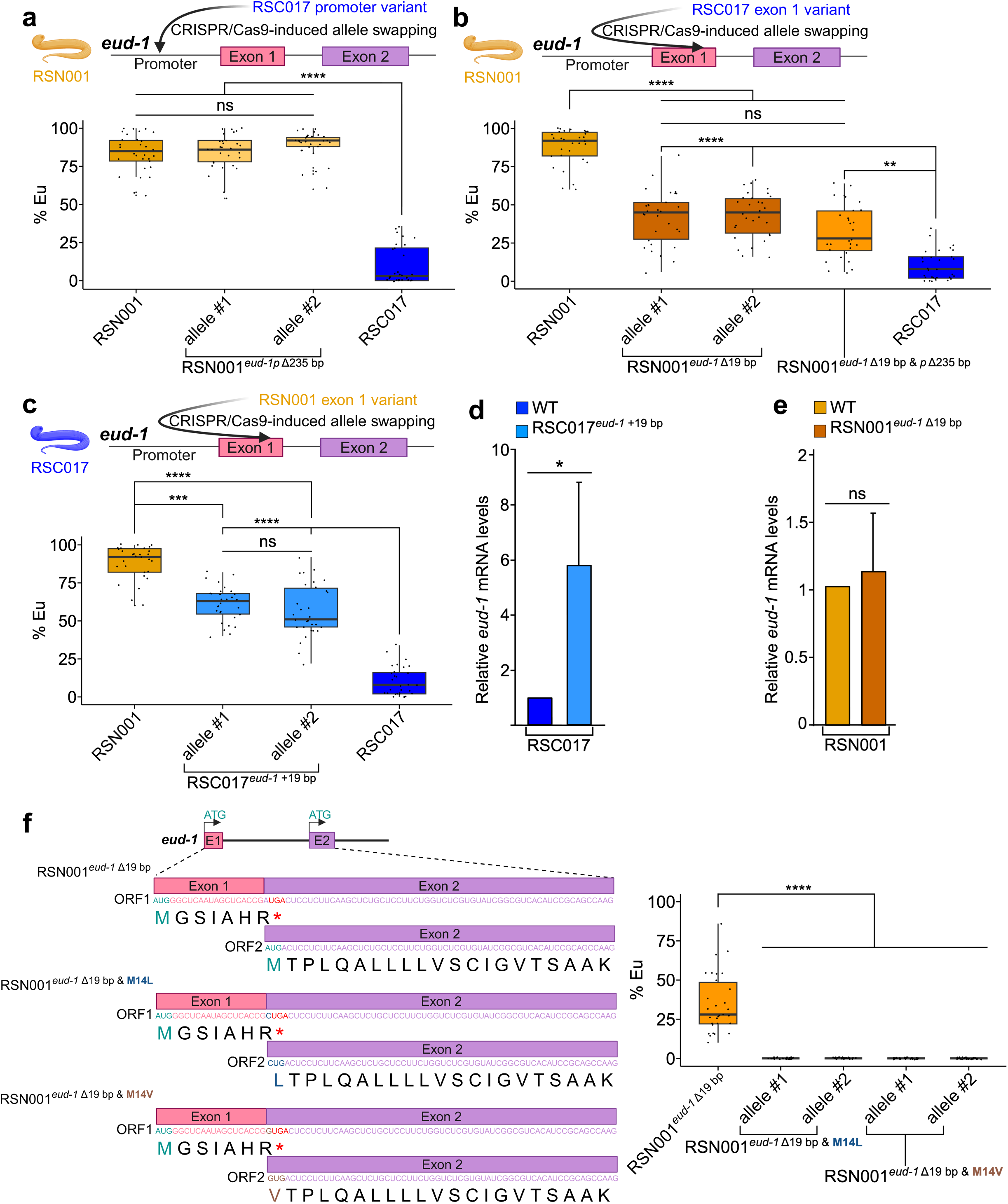
Elimination and reintroduction of the exonic 19 bp fragment partially alters mouth-form ratios. **a**–**c,** Percentage of Eu animals in RSN001 and RSC017 strains with genetic changes in the *eud-1* locus: (**a**) 235 bp deletion in the promoter region (*eud-1p* Δ235 bp) introduced into RSN001 by CRISPR engineering, (**b**) exonic 19 bp deletion (*eud-1* Δ19 bp) in RSN001, and (**c**) reintroduction of the exonic 19 bp (*eud-1* +19 bp) into the St-biased RSC017 strain. (*n* = 30 replicates). **d,e,** Determination of *eud-1* and *sult-8* mRNA levels by quantitative PCR (qPCR) following reverse transcription of RNA extracted from the different *P. pacificus* strains. *eud-1* mRNA levels were normalized to *sult-8* mRNA levels. Bars represent the mean value and error bars represent standard deviation (SD) (*n* = 3 biological replicates) **f,** Left, Gene scheme depicting *eud-1* exon 1 and exon 2 nucleotide and amino acid sequences with two in-frame methionines: M1 and M14 and the consequences of CRISPR-engineered genetic changes in nucleotide and amino acid sequences. Deletion of the 19 bp generates a premature termination codon (asterisk) at the beginning of exon 2, overlapping with the exon 2 *aug*. Point mutations resulting in the M14L and M14V amino acid substitutions remove the translation initiation site in exon 2 without disrupting the PTC. Right, quantification of Eu animals in these genetically altered RSN001 mutants (*n* = 30 replicates). **a**–**c,f,** Boxplots show the median (centre line), interquartile range (IQR; box), and whiskers extending to 1.5×IQR of mouth-form ratio. **a**–**f** Additional mutant information is shown in Supplementary Table 2. Statistical significance in **a**–**c,f** and **d,e** was assessed by Kruskal–Wallis test and the post-hoc Dunn’s test with a Holm–Bonferroni method and unpaired two-samples *t*-test, respectively. Significance levels are indicated as \*\*\*\**p* < 0.0001; \*\*\**p* < 0.001; \*\**p* < 0.01; * *p* < 0.05; ns (not significant) *p* > 0.05. Source data are provided as a Source Data file.

### Nonsense *eud-1* variants confer mouth-form plasticity

Next, we studied the potential role of the natural nonsense variant in exon 1 of *eud-1* (Fig. 3b,c). We had expected that the 19 bp deletion frameshift variant in RSC017 induces an all-St mouth-form phenotype similar to *eud-1* mutants isolated in forward genetic screens^13^, as it elicits premature translation termination in exon 2 (Fig. 2e). However, while RSC017 has a St-biased mouth-form ratio, around 10% of RSC017 worms still have the Eu morph (Fig. 1d). When we created CRISPR-engineered variants of this 19 bp fragment in the RSC017 and RSN001 backgrounds, we observed major shifts in mouth-form preferences. Specifically, a RSN001 worm population with the exact CRISPR-engineered 19 bp deletion (RSN001*^eud-1^* ^Δ19bp^) had less than 50% Eu mouth-form ratio (Fig. 3b). Similarly, the reintroduction of the 19 bp fragment into RSC017 (RSC017*^eud-1^* ^+19^ ^bp^), thereby recreating the reading frame of exon 1, increased the Eu mouth form from 10% to 50-60% in the population (Fig. 3c). These results indicate that the 19 bp deletion has a key role in the regulation of the mouth-form ratios in *P. pacificus* natural isolates. However, these shifts were not complete, which suggest that additional variants at the *eud-1* locus or other genetic loci contribute to the differential mouth-form preference between the two strains. RSN001 animals with both the 235 bp promoter deletion and the 19 bp deletion in exon 1 were still less than 50% Eu (Fig. 3b), confirming that the *eud-1* promoter deletion does not account for the effects observed in the mouth dimorphism.

One potential explanation for the absence of a ‘complete *eud-1* mutant phenotype’ would be that the formation of the Eu morph might be partially independent of EUD-1/sulfatase activity in these natural isolates. To test this hypothesis, we performed experiments with the steroid sulfatase inhibitor STX64^33,34^ that is known to prevent the development of the Eu morph in the *P. pacificus* PS312 reference strain^11^. We found that *P. pacificus* RSC017, RSN001 and the CRISPR-engineered mutant lines RSC017*^eud-1^* ^+19^ ^bp^ and RSN001*^eud-1^* ^Δ19bp^ were all-St when grown in the presence of STX64 (Suppl. Fig. 3). These findings suggest that all Eu animals seen in these cultures rely on EUD-1/sulfatase activity, and that the 19 bp deletion nonsense variant does not completely interfere with EUD-1 protein production.

### Variation in NMD strength accounts for changes in mouth-form phenotype

Nonsense-mediated mRNA decay (NMD) is an mRNA surveillance pathway that precludes translation of potentially harmful protein variants by affecting the stability of transcripts harbouring premature termination codons (PTCs)^35^. However, some nonsense transcripts escape NMD surveillance and undergo translation^36^. In particular, NMD is inefficient in detecting and targeting start-proximal PTCs^37–42^, such as the one observed in the *eud-1* nonsense variant (Fig. 2e). To shed light into the mechanism underlying the synthesis of EUD-1, we tested the possibility that strain-specific differences in *eud-1* nonsense mRNA levels result in the observed partial phenotypes. To determine the impact of the 19 bp deletion in *eud-1* expression, we measured *eud-1* mRNA levels in RSC017 and RSN001 animals with reintroduction (RSC017*^eud-1^* ^+19^ ^bp^) or deletion (RSN001*^eud-1^* ^Δ19bp^) of the 19 bp fragment, respectively. *eud-1* mRNA levels in RSC017 animals were approximately 6-fold increased following the correction of *eud-1* exon 1 sequence (Fig. 3d), suggesting that *eud-1* mRNA tends to be degraded by the NMD pathway following premature translation termination^43^ in the St-biased strain. This 6-fold increase in *eud-1* mRNA levels is accompanied by a 5-fold increase in the numbers of Eu animals in these lines (Fig. 3c). In contrast, in the Eu-biased RSN001 strain, *eud-1* mRNA levels were not altered upon deletion of the same 19 bp sequence (Fig. 3e). This finding suggests that in this strain, NMD is unable to induce the degradation of *eud-1* nonsense mRNAs. These findings are in agreement with other studies indicating that NMD activity is context-dependent^44^ and provide first evidence that NMD efficiency towards nonsense mRNAs with start-proximal PTCs differs among closely related wild isolates of *P. pacificus*.

### Alternative initiation of translation allows the synthesis of a functional EUD-1 sulfatase

Knowing that NMD is a translation-dependent process, we investigated possible mechanisms leading to the synthesis of a functional EUD-1 enzyme that results in the development of Eu animals in strains carrying the nonsense transcript variant in exon 1 (Suppl. Fig. 3). First, we considered whether inefficient termination of translation at the PTC via stop codon readthrough allows the ribosome to produce the EUD-1 sulfatase^40,41^. In this scenario, the ribosome continues translating and incorporates a non-cognate amino acid in the PTC codon^42^. The 19 bp deletion in *eud-1* exon 1 changes the frame of the ribosome; thus, stop codon readthrough would lead to the production of a protein with a distinct amino acid sequence unable to have the sulfatase activity required for the determination of the Eu morph (Fig. 2e). Therefore, our observations contradict the stop codon readthrough hypothesis, as Eu animals requiring EUD-1 activity are frequently observed in the different strains expressing the nonsense transcript (Fig. 3b,c and Suppl. Fig. 3).

We then evaluated the possibility that alternative translation initiation allows the synthesis of a functional EUD-1 enzyme. One striking feature of the *eud-1* mRNA is that it contains two potential alternative start *aug* in exon 1 (M1) and exon 2 (M14), respectively (Fig. 4a). Note that the second Met (M14) is still part of the predicted signal peptide of EUD-1 (Fig. 2c and 6b). One potential explanation for the absence of a ‘complete *eud-1* mutant phenotype’ (all-St) in RSC017 or RSN001*^eud-1^* ^Δ19bp^ would be that synthesis of EUD-1 utilises the *aug* at exon 2. The shortened signal peptide resulting from translational initiation in exon 2 would, according to SignalP 6.0 prediction^45,46^, not hinder effective translocation of the protein into the endoplasmic reticulum (ER) (Fig. 6b).

**Fig. 4.**
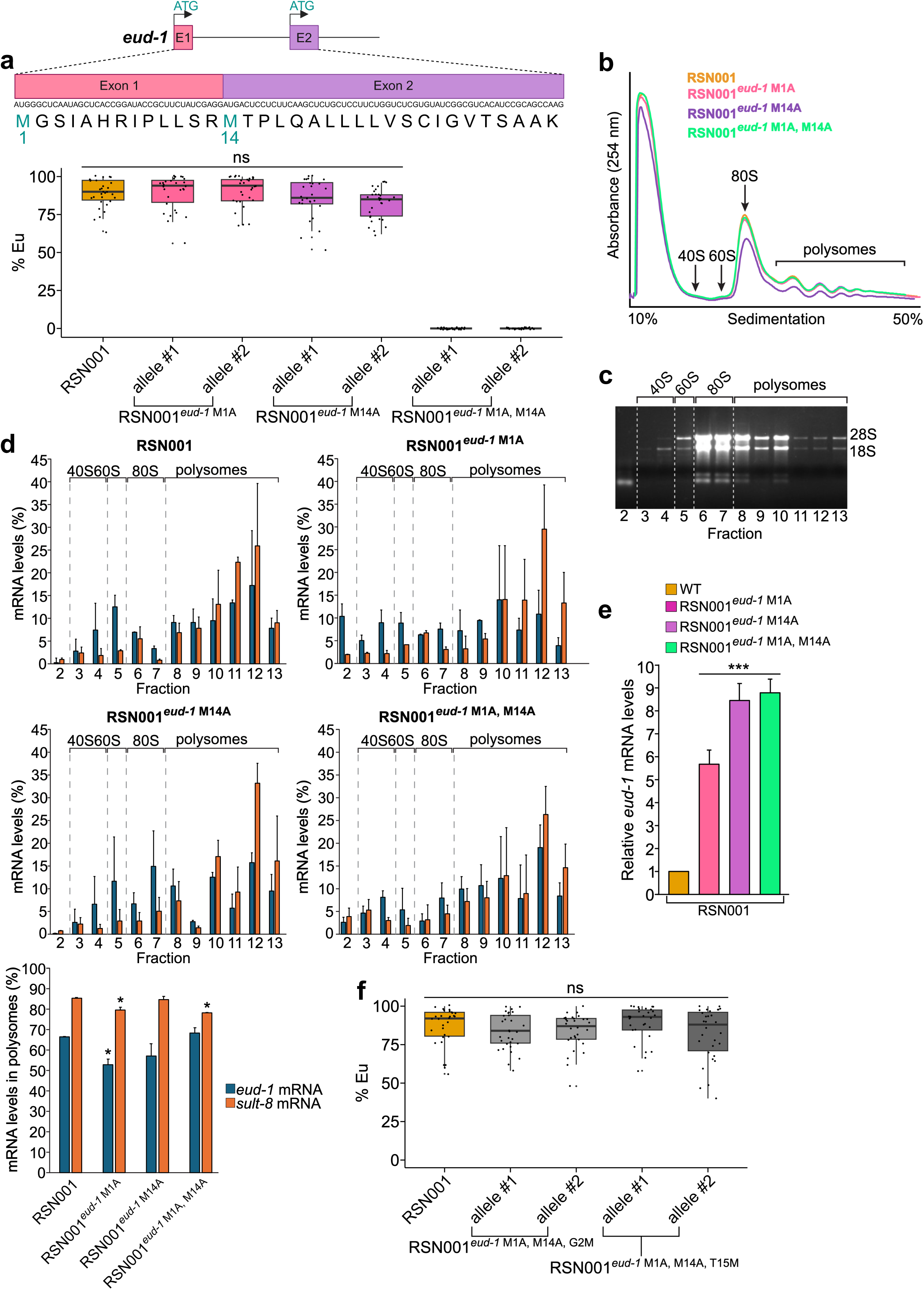
Alternative N-terminal EUD-1 proteoforms contribute to the Eu phenotype in RSN001. **a,** Schematic representation of *eud-1* exons 1 and 2 with nucleotide and amino acid sequences being depicted. The *eud-1* mRNA has two in-frame start sites: M1 or M14. The graph shows the percentage of Eu animals in wild type RSN001 animals and carrying single and double nucleotide substitutions disrupting the alternative translation start sites: M1A, M14A, and M1A M14A. Boxplots show the median (centre line), interquartile range (IQR; box), and whiskers extending to 1.5×IQR of mouth-form ratio (*n* = 30 replicates). **b,** UV absorbance profile at 254 nm of RSN001 wild type (orange) and mutants of *eud-1* alternative start codons (M1A, pink; M14A, purple; M1A, M14A, green) worm extracts following polysome sedimentation in a 10-50% sucrose gradient. Absorbance peaks at 254 nm representing free 40S and 60S subunits, 80S monosomes and polysomes are indicated. The polysome profiles from the different strains were similar without obvious changes in the translation status. **c,** Ethidium bromide staining of total RNA extracted from the sucrose fractions obtained with RSN001 wild type cell lysate. 18S and 28S rRNA bands are highlighted. **d,** Abundance profiles for *eud-1* (blue) and *sult-8* (orange) mRNAs across the density gradient in RSN001 wild type and mutants of *eud-1* alternative start codons (M1A, M14A and M1A, M14A) assessed using qPCR (top four panels). Estimation of the abundance of *eud-1* (blue) and *sult-8* (orange) mRNAs in polysomes (% RNA in fractions 8-13 of the density gradient) in the *P. pacificus* strains with different variants of *eud-1* alternative start codons (lower left panel). Bars represent the mean value and error bars represent SD (*n* = 3 biological replicates). **e,** Quantification of *eud-1* mRNA abundance in RSN001 wild type (WT) and in-frame methionine mutants using qPCR. *eud-1* mRNA levels were normalized to *sult-8* mRNA levels. Bars represent the mean value and error bars represent standard deviation (SD) (*n* = 3 biological replicates). **f,** Scoring of Eu animals in RSN001 following artificial reintroduction of a Met in the background of EUD-1 M1A, M14A mutants (**a**) at the second or fifteenth amino acid (M1A, M14A, G2M or M1A, M14A, T15M). Boxplots show the median (centre line), interquartile range (IQR; box), and whiskers extending to 1.5×IQR of mouth-form ratio (*n* = 30 replicates). **a,b,d**–**f** Additional mutant information is shown in Supplementary Table 2. Statistical significance in **a**,**f** and **d,e** was assessed by Kruskal–Wallis test and the post-hoc Dunn’s test with a Holm–Bonferroni method and one-way analysis of variance (ANOVA) and the post-hoc Dunnett’s test, respectively. Significance levels are indicated as \*\*\*\**p* < 0.0001; \*\*\**p* < 0.001; \*\**p* < 0.01; * *p* < 0.05; ns (not significant) *p* > 0.05. Source data are provided as a Source Data file.

To provide evidence for this hypothesis, we generated additional CRISPR-engineered *eud-1* variants. In the RSN001*^eud-1^* ^Δ19bp^ background, the 19 bp deletion shortens the nucleotide sequence between the second bases of the *cgg* (Arg 7) and *agg* (Arg 13) codons of exon 1, just before the Met 14 *aug* codon (Fig. 3f and 4a). The resulting reading frame is altered and an early *uga* stop codon that overlaps with, but does not destroy the coding potential of the alternative translation initiation site in exon 2, occurs in the potentially translated protein after amino acid position 8 (Arg 7 Stop 8, *cg**a ug**a*, Fig. 3f). We added an *a*>*c* or *a*>*g* nonsynonymous mutation in the third base of the *cg<u>a</u>* (Arg 7) codon. These nucleotide substitutions affect the start *aug* of exon 2 (resulting in *<u>c</u>ug* (M14L) or *gug* (M14V)) but do not remove the PTC (Fig. 3f). We found that CRISPR-engineered lines with either M14L or M14V substitution did not produce Eu animals and were all-St (Fig. 3f). Thus, when a nucleotide substitution modifies the alternative start codon in exon 2 (Fig. 3f), synthesis of a functional EUD-1/sulfatase in the nonsense *eud-1* mRNA is impaired, and animals are all-St. Mechanisms underlying the alternative use of start codons in mRNAs with start-proximal PTCs include leaky scanning, an event in which the ribosome scanning the mRNA misses the 5’ proximal *aug* triplet and proceeds until it recognises the next *aug*^47^, and/or reinitiation of translation following stop codon recognition^37–39^. As the efficiency of leaky scanning and reinitiation of translation is not 100%, but conditioned by multiple factors, both processes can explain the partial phenotype (50% Eu) of RSN001*^eud-1^* ^Δ19bp^ (Fig. 3b,f). Thus, variation in NMD efficiency together with regulation of translation initiation modulate the mouth dimorphism in *P. pacificus*.

### Mouth dimorphism is controlled by N-terminal EUD-1 proteoforms

Our results so far indicate that a EUD-1 protein with a shorter signal peptide is functionally relevant for the expression of the Eu phenotype in natural worm populations. To further explore the presence of alternative N-terminal proteoforms in the EUD-1 enzyme, we modified the *aug* codons in exons 1 and 2 to introduce M1A and M14A substitutions in the RSN001 background (lacking the PTC), respectively (Supplementary Tables 2 and 3). We generated two worm lines for each variant. None of the lines with RSN001*^eud-1^* ^M1A^ and RSN001*^eud-1^* ^M14A^ showed changes in the mouth-form ratio and animals remained largely Eu (Fig. 4a). One explanation for these findings is that translation can start at M1 or M14 and that both versions of the protein are functional. To support this notion, we measured the association of *eud-1* mRNA with ribosomes in the different RSN001 strains (wild type, M1A and M14A) using polysome profiling, following sucrose density gradient separation, and coupled with RT-qPCR (Fig. 4b–e). Similar to the RSN001 wild type version (66.4±5.5%), the majority of RSN001*^eud-1^* ^M1A^ (52.8±8.2%) and RSN001*^eud-1^* ^M14A^ (57.1±4.5%) *eud-1* mRNAs remained associated with polysomes indicating active translation (Fig. 4d). In agreement with the Eu phenotype in the RSN001 strains, translation from both start sites produces active N-terminal EUD-1 proteoforms. As the association of the single mutant *eud-1* mRNAs with ribosomes slightly decreases compared to the wild type version (Fig. 4d), our results also suggest that alternative translation initiation (as a result of leaky scanning) can modulate *eud-1* expression and the synthesis of N-terminal EUD-1 proteoforms.

Next, we created a double mutant with both *aug* codons being mutated (RSN001*^eud-1^* ^M1 A, M14A^). Strikingly, the RSN001*^eud-1^* ^M1A, M14A^ mutant strains have a fully penetrant mouth-form phenotype and result in all-St animals (Fig. 4a). This phenotype is similar to *eud-1* mutants originally isolated in the *P. pacificus* reference strain PS312^13,15^. These findings support the notion that M1 and M14 can both serve to initiate translation of N-terminal proteoforms of EUD-1. The all-St phenotype in the RSN001*^eud-1^* ^M1A, M14A^ animals was not caused by a reduction in *eud-1* mRNA levels, as the presence of single or double substitutions in the start *aug* codons increased *eud-1* mRNA abundance (Fig. 4e). We also measured the association of the RSN001*^eud-1^* ^M1A, M14A^ double mutant to ribosomes by polysome profiling. Curiously, the majority (68.3±3,5%) of mutant mRNAs were still associated with ribosomes (Fig. 4d). In the absence of M1 and M14, the most proximal *aug* codon is located after the signal peptide and at position 53 of the amino sequence of EUD-1 (Suppl. Fig. 4a). In the eventuality that the double mutant mRNA is still used for the translation of additional N-terminal variants of EUD-1, the lack of a signal peptide will result in the synthesis of an inactive enzyme as ER import is required for sulfatase maturation and activity^48^ (Suppl. Fig. 4b). In the presence of an inactive sulfatase, *P. pacificus* animals are exclusively St (Suppl. Fig. 3).

Finally, we used the RSN001*^eud-1^* ^M1A, M14A^ double mutant line and created a novel start *aug* at amino acid position 15 (Fig. 4f). We found that these RSN001*^eud-1^* ^M1A, M14A, T15M^ lines restored the high Eu ratio of the RSN001 wild type background (Fig. 4f). Similarly, a mutation resulting in an amino acid substitution G2M at position 2 in the M1A, M14A background (RSN001*^eud-1^* ^M1A, M14A, G2M^), restored the preferential Eu mouth-form phenotype (Fig. 4f). Taken together, these experiments indicate that alternative N-terminal EUD-1 proteoforms are produced in different *P. pacificus* wild isolates.

### Alternative N-terminal EUD-1 proteoforms are widespread among *P. pacificus* strains

We then analysed how conserved the *eud-1* two alternative *aug* start codons are in *P. pacificus* and the genus *Pristionchus*. We found that M1 and M14 are conserved among *P. pacificus* strains and closely related species, such as *P. exspectatus* and *P. arcanus,* also have the M1 and M14 arrangement (Suppl. Fig. 5a,b). To study if the observed alternative translation initiation of EUD-1 in RSN001 is functionally conserved, we generated similar mutants in the highly Eu PS312 strain. These mutant lines also contained a C-terminal ALFA tagged version of EUD-1 (Supplementary Table 2). We found that PS312*^eud-1^* ^M1A^ and PS312*^eud-1^* ^M14A^ animals maintain the Eu-biased mouth-form ratio, whereas PS312*^eud-1^* ^M1A, M14A^ worms are all-St (Fig. 5a). In agreement with mouth morphology, using anti-ALFA immunostaining we observed that EUD-1 was produced in the pharyngeal (head) sensory neurons of PS312 wild type, PS312*^eud-1^* ^M1A^ and PS312*^eud-1^* ^M14A^ animals, but not in the same corresponding cells of PS312*^eud-1^* ^M1A, M14A^ worms (Fig. 5b). These results indicate that also in the *P. pacificus* PS312 strain, the M1 and M14 *aug* codons can be alternatively used in the translation of *eud-1* mRNA that produces a functional enzyme. Together, these findings suggest that alternative N-terminal proteoforms are widespread and conserved in different *P. pacificus* clades.

**Fig. 5.**
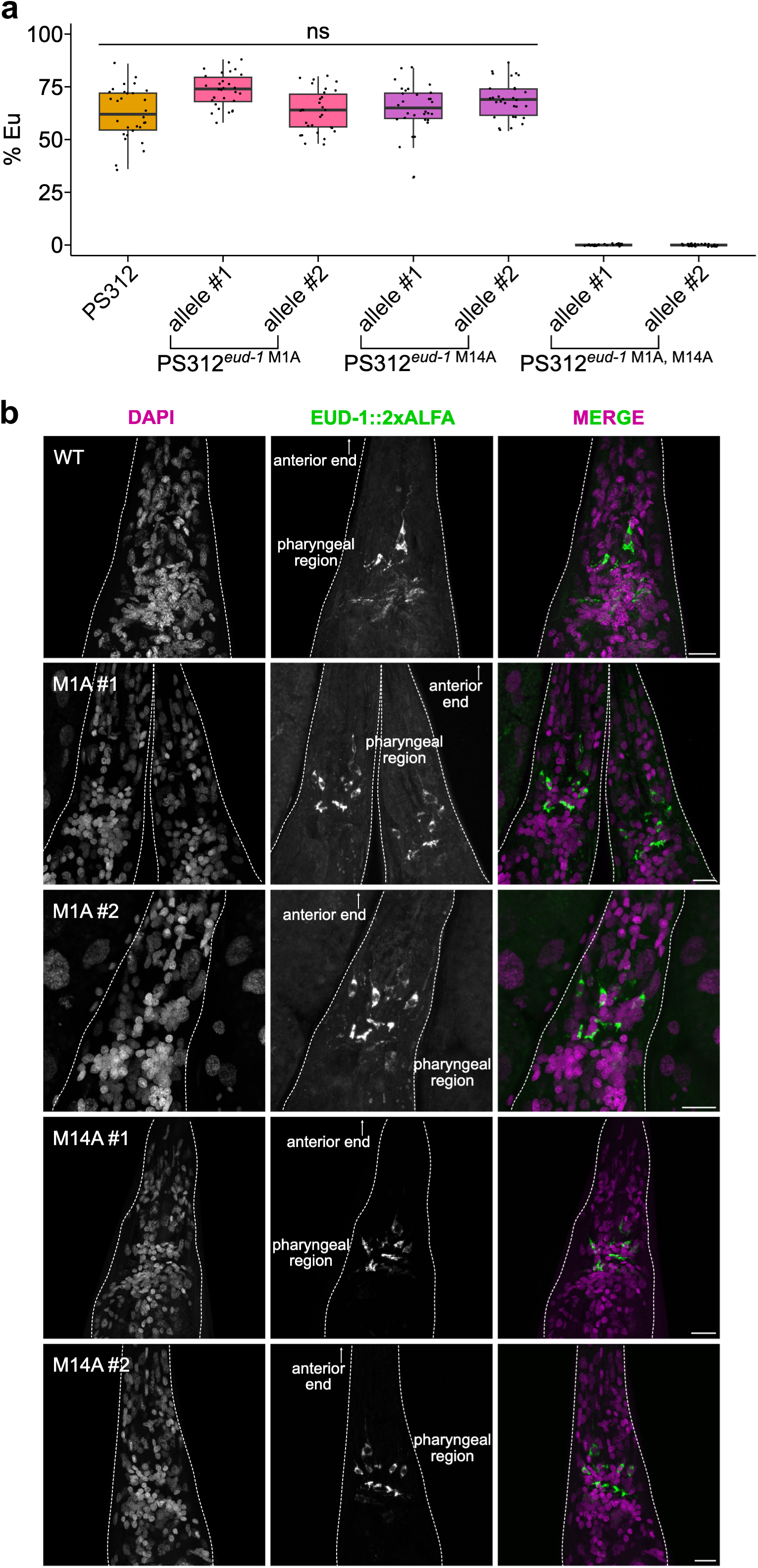
N-terminal EUD-1 proteoforms are synthesised in pharyngeal (head) sensory neurons of *P. pacificus* animals. **a,** Graph depicts the percentage of Eu animals in wild type and *eud-1* CRISPR-engineered (M1A, M14A, M1A M14A) *P. pacificus* PS312 strain. Boxplots show the median (centre line), interquartile range (IQR; box), and whiskers extending to 1.5×IQR of mouth-form ratio (*n* = 30 replicates). Statistical significance was assessed by Kruskal–Wallis test and the post-hoc Dunn’s test with a Holm–Bonferroni method. Significance levels are indicated as \*\*\*\**p* < 0.0001; \*\*\**p* < 0.001; \*\**p* < 0.01; * *p* < 0.05; ns (not significant) *p* > 0.05. Source data are provided as a Source Data file. **b,** Representative fluorescent confocal images of the pharyngeal head region of fixed adult *P. pacificus* PS312 wild type (WT), and both alleles (#1 and #2) of EUD-1 M1A and M14A mutant animals. These strains have a C-terminal ALFA tagged version of EUD-1. All animals were stained with anti-ALFA nanobody coupled with AZDye568 fluorophore. Merged pictures depict the anti-ALFA signal in green and the nuclei (DAPI) in purple. Arrows indicate the anterior end of the animals. Scale bars: 10 µm. Additional mutant information is shown in Supplementary Table 2.

### Alternative N-terminal proteoforms are common in eukaryotic sulfatases

Finally, we wanted to know if alternative N-terminal proteoforms are specific to EUD-1 or if such forms are also found in sulfatases of other nematodes or even more distantly related animals and humans. The *eud-1* gene arose from a recent duplication of *sul-2* in the genus *Pristionchus* and is one of three *sul-2* paralogs (*eud-1*, *PPA06135* and *PPA21290*) in *P. pacificus*^13^. Curiously, the signal peptide (SP) sequence of PPA21290 also contains multiple Met residues (M1, M4, M12, M13 and M19) (Fig. 6a,c). *In silico* analysis of PPA21290 N-terminal proteoforms at these different residues indicates that with the exception of the M19, initiation of translation at the different Met residues allows the synthesis of this SUL-2 enzyme with functional SP sequences (Fig. 6c). Although unknown to date, this observation suggests that alternative N-terminal proteoforms of PPA21290 might also occur in *P. pacificus*.

**Fig. 6.**
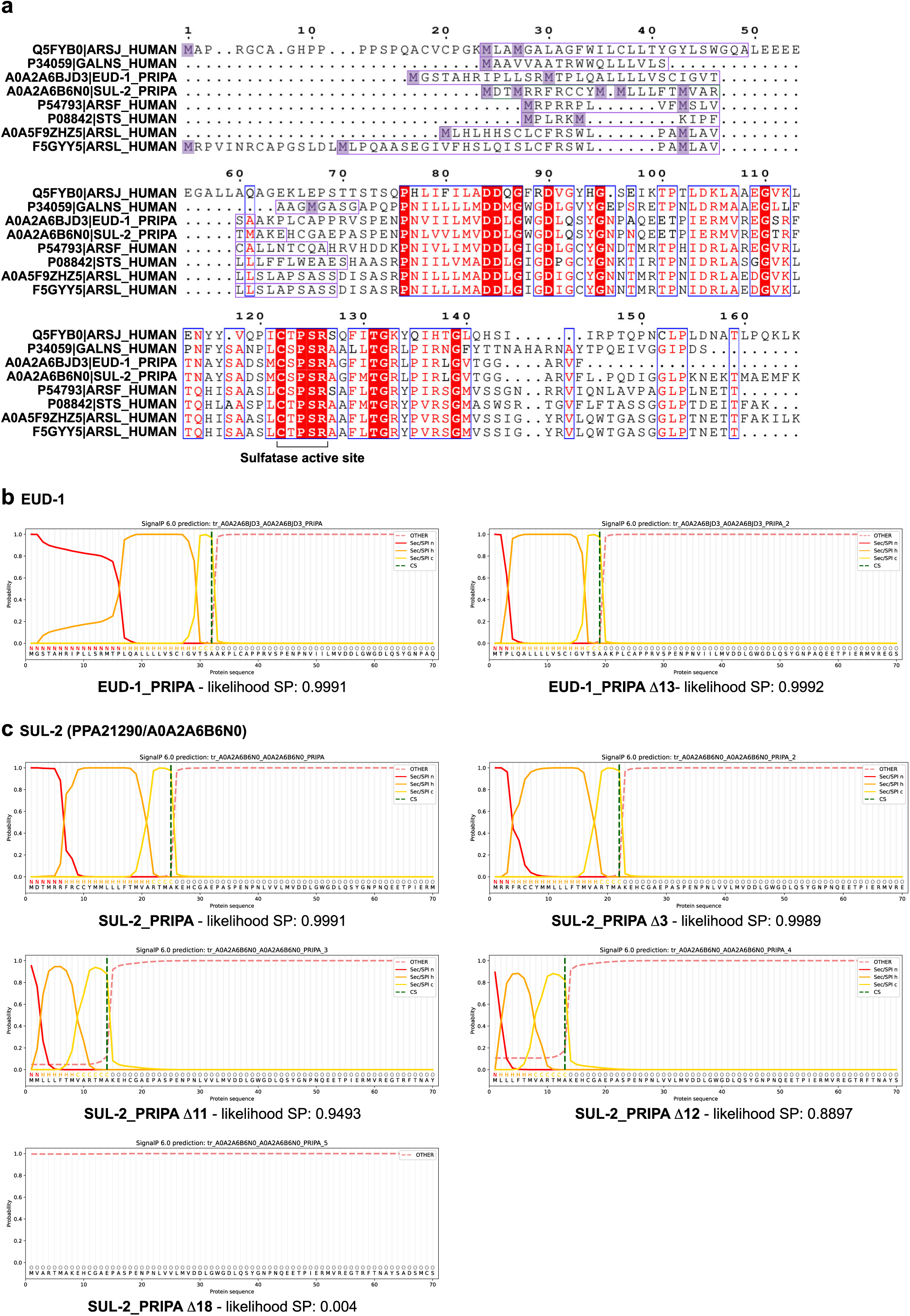
Alternative N-terminal EUD-1 proteoforms are common in *P. pacificus* sulfatases. **a,** Sequence alignment of the N-terminal regions of the human arylsulfatases ARSJ (Q5FYB0), GALNS (P34059, N-acetylgalactosamine-6-sulfatase), ARSF (P54793), STS (P08842, steroid sulfatase), and ARSL long (F5GYY5) and short (A0A5F9ZHZ5) isoforms, and *P. pacificus* (PRIPA) SUL-2 sulfatases EUD-1 (A0A2ABJD3) and PPA21290 (SUL-2, A0A2A6B6N0) performed with ESPript 3.0. Identical residues are identified in red columns and printed in white whereas similar residues are shown in red. The signal peptide (SP) amino acid sequence for each sulfatase is highlighted by a purple box. Methionine (M) residues are indicated in purple squares. The active site of the enzymes is also indicated. **b,c,** Prediction of signal peptides (n-, h- and c-regions) and cleavage sites (CS) using SignalP 6.0 in the N-terminal proteoforms of EUD-1 (**b**) and PPA21290 (**c**) SUL-2 nematode sulfatases. Δx denotes the predicted proteoform starting from Mx+1. Likelihood values for each prediction are denoted below each graph.

Nematode SUL-2 sulfatases belong to the arylsulfatase enzyme family with activity towards steroid hormones^49^. Interestingly, in human cells, the expression of the steroid sulfatase (STS) is modulated by different promoters mediating the synthesis of transcripts with alternative first exons. The resulting STS enzymes display functional variations of the SP sequence (Fig. 6a, 7a) and are expressed in a cell-type/tissue-specific manner^50–52^. Human STS is not the only eukaryotic sulfatase with reported or putative N-terminal variants. Indeed, a survey at the SP sequences of human sulfatases for the presence of multiple Met residues also identified the arylsulfatases ARSJ (Q5FYB0), GALNS (P34059), ARSF (P54793) and short and long isoforms of ARSL (A0A5F9ZHZ5 and F5GYY5) (Fig. 6a, 7b–f). Such as in the case of human STS or EUD-1, usage of alternative *aug* codons will produce variants of the ARSJ and ARSL enzymes with functional SPs (Fig. 7b,e,f). Taken together, these observations suggest that alternative N-terminal proteoforms are common in eukaryotic sulfatases. However, despite the roles of sulfatases in development and diseases, the detection, roles and mechanisms underlying the expression of N-terminal variants of these enzymes remain understudied.

**Fig. 7.**
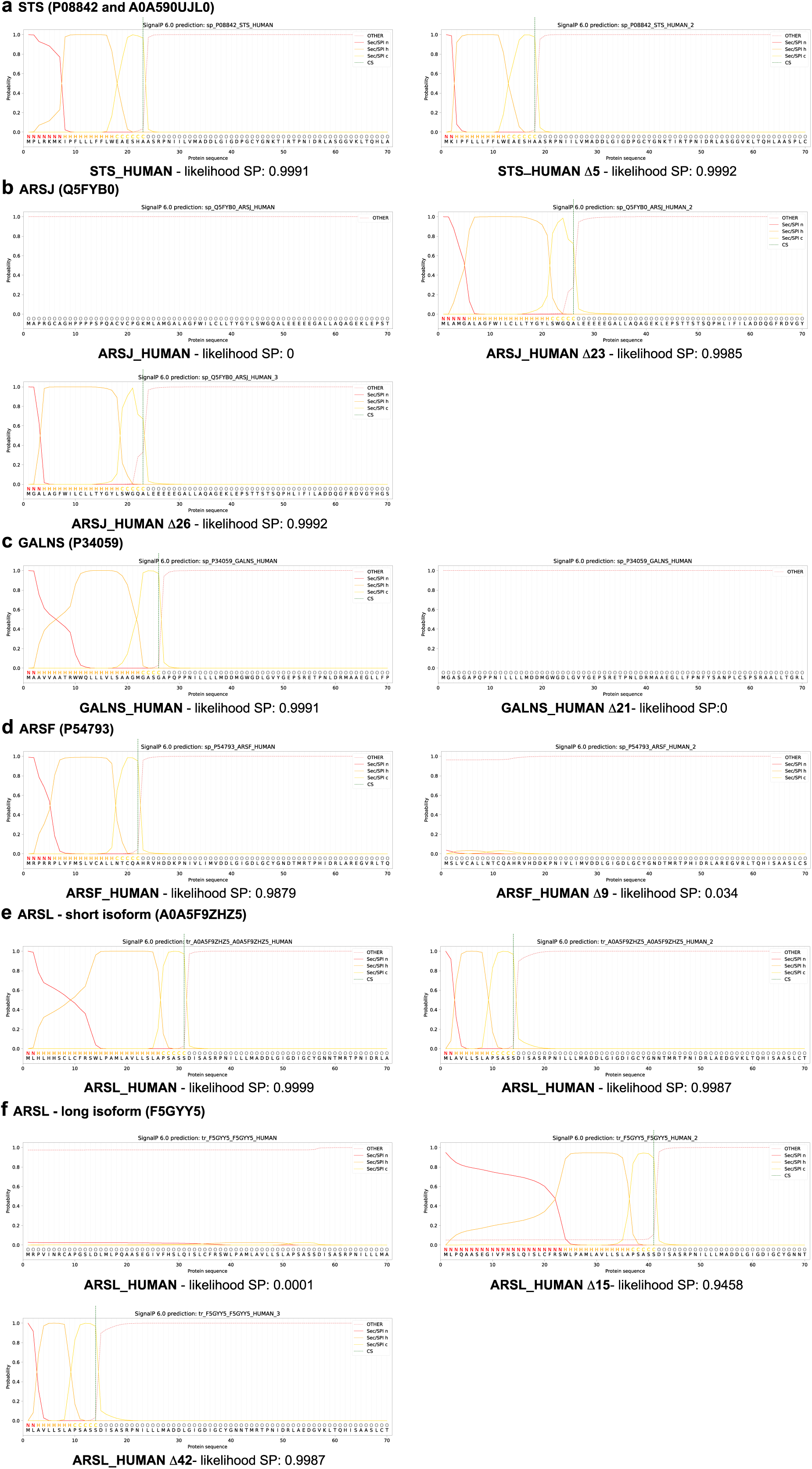
Alternative N-terminal EUD-1 proteoforms are common in eukaryotic sulfatases. **a-f,** Analysis of SP and cleavage sites following alternative start site usage in human sulfatases using SignalP 6.0. Proteins are as follow: (**a**) steroid sulfatase (STS, P08842, A0A590UJL0), (**b**) ARSJ (Q5FYB0), (**c**) GALNS (P34059, N-acetylgalactosamine-6-sulfatase), (**d**) ARSF (P54793), (**e**) ARSL short isoform (A0A5F9ZHZ5) and (**f**) ARSL long isoform (F5GYY5). Δx denotes the predicted proteoform starting from Mx+1. Likelihood values for each prediction are denoted below each graph.

## Discussion

This work provides a molecular explanation for variation in mouth-form bias in natural isolates of the nematode plasticity model *P. pacificus* carrying nonsense variants of the *eud-1* developmental switch locus, thereby linking evolutionary and ecological aspects of developmental plasticity to molecular mechanisms. Our findings are derived from large scale population surveys, organismal-specific genetic manipulations and multiple biochemical assays. We provide evidence for variability of NMD in natural populations and distinct translational control mechanisms regulating the synthesis of variants of a sulfatase with a key role in plasticity.

We observed that NMD strength differs among *P. pacificus* natural populations, as the abundance of the *eud-1* Δ19 bp nonsense mRNA differed from its wild type counterpart in RSC017 animals, but not in the RSN001 animals. Variability in NMD efficiency is observed in animals, plants and fungi across environmental and developmental conditions, genetic background, cell type and individuals^53–55^.This variation in NMD efficiency results in phenotypic variability and different degrees of expressivity of a nonsense *eud-1* variant. Our study supports the role of NMD in facilitating adaptive evolution by modulating the expression of cryptic genetic variation in response to environmental stimuli^44^. Therefore, we speculate that mechanisms that turn on and off the expression of plastic traits will facilitate evolutionary capacitance by increasing the potential of an organism to adapt and evolve when faced with novel environmental or genetic constraints. Although the mechanisms linked to variations in NMD robustness in *P. pacificus* wild isolates require further investigation, this collection of strains isolated over several years can provide valuable information on the molecular complexities of the NMD pathway and holds promise for the identification of novel therapeutic strategies to counter genetic disorders caused by nonsense mutations. In addition, our results provide a molecular explanation for the synthesis of a functional sulfatase from the nonsense *eud-1* transcript. CRISPR-targeted disruption of in-frame start codons and biochemical assays indicate that the presence of alternative *aug* in *eud-1* transcript mediates the synthesis of functional N-terminal proteoforms of the sulfatase by leaky scanning and/or translation reinitiation.

The N-terminal region of sulfatases performs a crucial function in enzyme activity. It contains the signal peptide that directs sulfatases to the ER where they undergo a co-translation modification essential for activity^48^. Despite the restricted number of studies addressing N-terminal proteoforms in sulfatases, alternative promoter usage has been proposed to elicit the synthesis of STS enzymes with functional variations of the signal peptide in a tissue-dependent fashion^50–52,56^. Our analysis of nematode and human sulfatases N-terminal sequences highlights the presence of in-frame methionines in multiple enzymes and the possibility of alternative translation start sites to modulate signal peptide structure or function, adding a new and previously unrecognisable layer to the regulation of enzymes with crucial role in hormone regulation, modulation of signal pathways, and cellular degradation^57^.

In conclusion, using a survey englobing ecological, genetic and biochemical studies on a nematode phenotypic polyphenism, we uncover molecular mechanisms with vast implications in animal biology and evolution.

## Methods

### Beetle collection and nematode isolation

Beetles of the genus *Adoretus* Dejean, 1833 were hand collected from Parc du Colorado, La Réunion^58^ (Fig. 1c). They were brought alive to the Max Planck Institute for Biology Tübingen to be sacrificed on nematode growth medium (NGM) plates seeded with *Escherichia coli* OP50. A cocktail of *E. coli* and proliferating native beetle carcass bacteria induce the emergence of nematode dauers. *P. pacificus* isolates were established as a single gravid isogenic hermaphrodite using morphological identification^59^.

### *P. pacificus* population genomics

A genome-wide phylogeny of diverse *P. pacificus* strains was generated by sequencing the genomes of newly isolated strains and integrating them into the previously published population genomic data sets for *P. pacificus*^23,26,27^ (Suppl. Fig. 1). To this end, the whole genome sequencing datasets were aligned against the genome assembly of the *P. pacificus* reference strain PS312 (version El Paco) with the software BWA (version 0.7.17-r1188, with option mem)^27,60^. The resulting alignment files were then genotyped by the samtools mpileup (version 0.1.18 r982:295) and bcftools (version 0.1.17-dec r973:277) at a set of candidate positions that were previously found to be variable across different *P. pacificus* natural isolates^23,26,27,61^. Variable positions that could be genotyped in all strains with a quality score of at least 20 were concatenated into an artificial multiple sequence alignment that was taken as an input by the phylogenetic R package phangorn in order to calculate a neighbour-joining tree based on Hamming distances between strains^62^.

### *P. pacificus* mouth-form phenotype assays

The modular stereomicroscope ZEISS SteREO Discovery.V12 was used to determine the mouth form of adult *P. pacificus* hermaphrodites. Overall stomatal differences were used to determine the mouth-form phenotype. The Eu individuals have a wider and more sclerotised stoma than St animals^13^ (Fig. 1a).

All mouth-form assays were performed in standard laboratory conditions with *E. coli* OP50 as food source and nematode cultures grown at 20°C. For each genotype, an individual J4 was picked onto an NGM plate seeded with 300 µl of *E. coli*. The mouth-form phenotyping was done for their progeny six days later. For *P. pacificus* natural isolates, phenotyping was conducted on five technical replicates (plates) over three independent experiments (*n* = 15). Depending on the genotype’s fecundity, thirty to fifty animals were phenotyped per plate to determine the Eu mouth-form ratio (Fig. 1d). For the rest of the genotypes, mouth-form assays were carried out with three biological replicates (10 plates per biological; *n* = 30). The proportion of Eu phenotype was determined from fifty animals per plate.

### Recombinant inbred line (RIL) analysis

Self-sperm-depleted hermaphrodites and males were needed to generate RILs. Late J4 hermaphrodites were picked and transferred daily to a new plate for four days at which point self-sperm ran out. To induce males for crosses, two spontaneous males of each strain were crossed with five self-sperm depleted hermaphrodites of the corresponding genotype on a 100-µl *E. coli* OP50 lawn.

To generate a panel of RILs, P0 males of *P. pacificus* RSC017 and self-sperm-depleted hermaphrodites of *P. pacificus* RSN001 were crossed at a ratio of 2:5. A reciprocal cross was also set up to balance the panel^63^. Parents were removed after ∼16 h. A day later, F1 J4-stage hermaphrodites were singled out and left to lay F2 eggs before worm lysis. Sanger sequencing was used to confirm the heterozygosity using a single nucleotide polymorphism (SNP) (*g* in RSN001 > *a* in RSC017) at ChrI:3434406. The primer pair is provided in Supplementary Table 3. All tested F1 animals were heterozygous and 80 F2 individual J4 hermaphrodites were isolated per cross. A total of 160 RILs was transferred as individual J4 hermaphrodites until near homozygosity was reached at F12 (Fig. 2a). These lines were subsequently subjected to mouth-form phenotyping as described above. Eu mouth-form ratio was determined from thirty individuals per plate. Three replicates per RIL were assayed (*n* = 3) (Suppl. Fig. 2).

RILs were genotyped using in-house Tn5-transposase-based single worm whole genome sequencing protocol as previously described^64^. Briefly, a single RIL individual was lysed in 20 µl of worm lysis buffer. One fifth of the lysate was used for DNA cleanup by Sera-Mag Speedbead carboxylate-modified [E3] magnetic particles (Cytiva Cat. No. 65152105050250) at a ratio of 1:9. The purified DNA sample was then subject to tagmentation by in-house-produced Tn5 transposase (a gift from the Weichenrieder lab, MPI for Biology, Tübingen). This step was followed by a specific i5-and-i7-barcoded 14-cycle PCR. The resultant PCR product was size-selected for fragments between 300 and 600 bp. Afterwards, DNA concentration and size were determined using Qubit (Thermo Fisher) and Bioanalyzer (Agilent), respectively. Finally, a total of 141 RILs’ DNA samples were pooled and sequenced at Novogene.

### Genome assembly and gene annotation of the RSC017 strain

To establish genomic resources for the *P. pacificus* strain RSC017, we sequenced and assembled its genome following recently established protocols^65^. In short, a PacBio library was generated, sequenced and subsequently assembled into a raw assembly by the software Canu (version 1.4)^66^. This assembly was scaffolded by the software RagTag (version 2.1.0) with the help of the chromosome-scale *P. pacificus* assembly for the strain PS312 (version El Paco)^27,67^. Protein-coding genes were annotated by the software PPCAC which incorporates strain-specific RNA-seq data and the community-curated *P. pacificus* gene annotations (version El Paco annotations 3)^68,69^.

### QTL mapping

To find associations between genotype and phenotype of the RILs, we followed a similar protocol as described previously^28^. In summary, whole genome sequencing data of the RILs were aligned against the RSC017 reference genome using BWA mem and genotyped at variable positions by samtools mpileup bcftools^27,61^. Subsequently, the logarithm of the odds (LOD) scores for the association between genotype and phenotype (high or low mouth-form ratio; Suppl. Fig. 2) were calculated at 50,000 randomly selected variant positions as the negative log_10_ of the *p*-value of a Fisher’s exact-test. A LOD score above six was considered as a significant association (Bonferroni correction).

### Analysis of QTL gene candidate sequence variants

The software Integrative Genomics Viewer (IGV; version 2.16.2) was used to visually analyse the *eud-1* locus sequence variants between the RIL parental strains, RSN001 and RSC017, and the rest of strains tested for mouth-form in this study^70^ (Fig. 1d and 2d and Supplementary Table 1). *P. pacificus* PS312 genome and gene annotations were used as the reference to which bam files of natural isolates were aligned. The observed sequence variants in the *eud-1* promoter and exon 1 between RSN001 and RSC017 were confirmed by Sanger sequencing using primers provided in Supplementary Table 3.

### CRISPR/Cas9 mutagenesis

The Eppendorf microinjection system was used for gonadal delivery of CRISPR/Cas9-sgRNA mix into well-fed late-J4 hermaphrodites following a well-established protocol in *P. pacificus*^71^. An sgRNA (single guide RNA) was prepared by forming a duplex between a target-specific CRISPR RNA (crRNA) and a trans-activating crRNA (tracrRNA) at an amount of 5 µl each. This mix was incubated at 95°C for 5 min followed by another 5-min incubation at room temperature (RT). crRNAs (Supplementary Table 3) and tracrRNA (Cat. No. 1072534) were synthesised by Integrated DNA Technologies (IDT). Target-specific crRNAs are 20-nucleotide long and were designed to be immediately upstream of protospacer adjacent motifs (PAMs). A CRISPR/Cas9-sgRNA mix was prepared by mixing 5 µl of sgRNA with 1 µl of Cas9 endonuclease (IDT Cat. No. 1081058). This solution was incubated for 5 min at RT and 2 µl of a specific repair template synthesised by IDT (Supplementary Table 3) were added. A final concentration of approximately 25 ng/µl of *Ppa-eft-3p*::RFP was added to the mix as a coinjection marker before 1xTE (Promega Cat. No V6231) was added to bring the total volume to 20 µl. Finally, this solution was centrifuged at maximum speed at 4°C for 10 min to remove any precipitates before injection. The final concentration of specific sgRNA, Cas9, *Ppa-eft-3p*::RFP and repair template is 12.5 µM, 0.5 µg/µl, 25 ng/µl and 10 µM, respectively.

Injected P0 were individually recovered and let lay eggs on a 50-µl *E. coli* OP50 lawn for ∼16 h before getting sacrificed. F1 progeny were screened for red fluorescence, an indication of successful gonadal injection. From a positive plate, 16 hermaphrodites were randomly singled out, let lay F2 eggs and Sanger sequenced. F1 heterozygotes with the correct DNA editing (Supplementary Table 3) as shown by sequence alignment to the wild type sequence using the MAFFT online service^72^ were further singled out at F2 to reach homozygosity. Two independent alleles were used for further assays and frozen in liquid nitrogen for long-term storage (Supplementary Table 2).

### Steroid sulfatase inhibitor STX64 experiments

To assay the activity of *P. pacificus* sulfatase EUD-1, the steroid sulfatase inhibitor STX64 (Sigma-Aldrich Cat. No. S1950) was used following a previous study^11^. Ten hermaphrodites of each *P. pacificus* genotype (Suppl. Fig. 3) were picked onto an NGM plate seeded with a 300-µl-*E. coli*-lawn. Five plates were prepared per genotype. After five days, population synchronisation was done from the egg stage using the routine bleaching protocol in nematodes^73^. These eggs were plated onto freshly prepared STX64 or the control DMSO (Sigma-Aldrich Cat. No. D8418) plates. These plates were prepared by seeding with 200 µl of OP50 + 10 µl of STX64 (1mg/ml in DMSO) or 10 µl of DMSO for a final concentration of ∼1 µg/ml and left to dry in the dark before the assay. After 72 h at 20°C, the mouth-form phenotype of 50 bleach-synchronised adult hermaphrodites was scored. The assays were performed in three biological replicates with 10 plates per replicate (*n* = 30).

### RNA extraction, reverse transcription (RT) and quantitative PCR (qPCR)

*P. pacificus* mutants and corresponding wild type animals (Supplementary Table 2) were harvested from three 6 cm Petri dishes of mixed stage cultures with M9 medium. Total RNA was isolated from pelleted animals using 1 ml of TRIzol Reagent (Invitrogen Cat. No. 15596018), according to the manufacturer’s instructions, with minor changes. Before phase separation, three cycles of flash freezing in liquid nitrogen and thawing at 37°C with vigorous shaking were conducted on the TRIzol resuspended animals to achieve complete worm lysis. DNase treatment and RNA purification were performed with the RNA Clean and Concentrator Kit-25 (Zymo Research Cat. No. R1018). 1 µg of RNA was reverse transcribed using the manufacturer protocol of SuperScript III Reverse Transcriptase (Invitrogen Cat. No. 18080044) in a reaction containing additional 2.5 µM of oligo (dT)_20_ (Jena Bioscience Cat. No. PM-304), 0.5 mM dNTPs, 40 U RNaseOUT recombinant ribonuclease inhibitor (Invitrogen Cat. No. 10777019), and 200 U of the reverse transcriptase. The qPCR reactions were performed with Luna Universal qPCR Master Mix (New England Biolabs Cat. No. M3003), with 0.25 µM of each primer and 1 and 2 µl of the cDNA for *sult-8* and *eud-1* reactions, respectively. Transcript-specific qPCR primers are listed in Supplementary Table 3. Normalised transcript expression ratios from three independent experiments (*n* = 3) were determined using the Livak-Schmittgen method^74^.

### Polysome profiling

To obtain the amount of worm material required for polysome profiling assays in triplicate, seven young hermaphroditic animals of each worm strain (Supplementary Table 2) were picked onto 40 plates 6-cm diameter NGM plates with *E. coli* OP50 as food source.

Four to five days later and before the worms run out of food, plates were chunked into small agar cubes with worms, and each cube was transferred onto 100-150 10-cm plates with *E. coli* OP50. Three to four days later, non-starved animals from the mixed stage cultures were collected with M9 buffer into Falcon tubes. Following three gentle washes in M9 and filtration on a 5 µm Nylon net filter (Millipore Cat. No. NY0504700) to remove remaining bacteria, and one wash in hypotonic buffer (10 mM Tris-HCl pH 7.9, 2.5 mM MgCl_2_), worms were flash frozen in liquid nitrogen and stored at −80°C. In total, 1-2 g of frozen worm popcorns were collected per strain. Worm popcorns were crushed with a pestle to fine powder on a mortar pre-cooled with liquid nitrogen. Worm powder was distributed between different microfuge tubes. Per polysome profiling assay, 400-500 µl (w/v) of worm powder was thawed and lysed in 1x vol of hypotonic lysis buffer containing 0.1 mM of cycloheximide (Carl ROTH Cat. No. 8682), 1 mM DTT, 0.5% Nonidet P40 (MP Biomedicals Cat. No. RIST1315), 1% sodium deoxycholate monohydrate (DOC) (Sigma Aldrich Cat. No. D6750) and 2% polyoxyethylene-10-tridecylether (PET) (Sigma Aldrich Cat. No. P2393) on ice. Following 40 strokes with an Eppendorf micropestle, worm lysates were kept on ice for 10 min. Worm debris was removed by a 10 min centrifugation at 12 000 *g* at 4°C. The soluble lysate was transferred to a fresh tube and absorbance was measured at 260 nm. 70 U A260 of worm lysate were layered on a linear 10-50% (w/v) sucrose gradient, prepared using Gradient Master 107ip (Biocomp) in gradient buffer (10 mM Tris-HCl pH 7.9, 2.5 mM MgCl2, 80 U units RNase OUT (Invitrogen Cat. No. 10777019)/RiboLock RNase inhibitor (Thermo Scientific Cat. No. EO0382), 0.1 mM cycloheximide. Centrifugation was carried in a Beckman ultracentrifuge using a SW55Ti at 45 000 rpm, for 45 min at 4°C. Polysome fractions were collected using the TRIAX Gradient Fractionation System Model FC-2 (Biocomp) and polysomes profiles were monitored by absorbance of light at 254 nm.

Samples for RNA isolation were first digested with proteinase K (Sigma Aldrich Cat. No. 1.24568, 1% of the sample volume; 100 mg/ml in 50 mM Tris-HCl pH=8.1, 10 mM CaCl_2_ buffer) at 37°C for 45 min and shaking at 400 rpm. The digested sucrose fractions were mixed with 1 volume of Phenol:Chloroform:Isoamyl alcohol (PanReac AppliChem, 25:24:1, v/v), vortexed and spun down 5 min at 20,000 *g* at 4°C. Supernatants were transferred into 3 volumes of 100% ethanol, 0.1 volumes of 3M NaOAc pH=5.2 and 1 µl of GlycoBlue (Ambion Cat. No. AM9515), and precipitated at −20°C. Samples were pelleted for 30 min at 20,000 *g* and 4°C, washed once with 100% ethanol and another time with 70% ethanol, dried and resuspended in 15 µl H_2_O. Fractions were reverse transcribed and analysed by RT-qPCR.

### Statistical tests

Statistical tests were performed using R language-based packages (R version 4.3.1)^75^. Homogeneity of variances and normal distribution were tested using Levene’s and Shapiro–Wilk tests, respectively. *eud-1* mRNA and polysome profiling data met the assumptions for parametric tests, i.e. the data are normally distributed and variances are homogenous. The unpaired two-samples *t*-test was used to analyse *eud-1* mRNA levels in wild type and 19 bp deletion or insertion mutants (Fig. 3d,e). The one-way analysis of variance (ANOVA) and the post-hoc Dunnett’s test were used for statistical tests of *eud-1* and *sult-8* mRNA levels in the polysome fractions (Fig. 4d,e). Mouth-form ratio data did not meet the parametric test assumptions, thus the non-parametric Kruskal–Wallis test and the post-hoc Dunn’s test with a Holm–Bonferroni method were used to test each pair (Fig. 3a–c,f, 4a,f and 5a). Significance levels were indicated as \*\*\*\**p* < 0.0001; \*\*\**p* < 0.001; \*\**p* < 0.01; * *p* < 0.05; ns (not significant) *p* > 0.05.

### Immunofluorescence

ALFA-tagged EUD-1 protein immunofluorescent staining was performed according to the protocol previously described^76^. Briefly, *P. pacificus* PS312 wild type and EUD-1 M-to-A mutants from mixed stage cultures were collected from ten 6-cm NGM plates, filtered on a 5-µm Nylon net filter (Millipore Cat. No. NY0504700) to remove remaining bacteria, and fixed with a 4% para-formaldehyde, 1x PBS solution overnight at room temperature on a tube rotator. Fixed animals were washed thrice with 0.5% Triton X-100, 1x PBS (washing solution), and incubated in 5% β-mercaptoethanol, 1% Triton X-100, 0.1M Tris-HCl pH 7.4 solution overnight at 37°C and shaking (300 rpm). Worm cuticle digestion was performed at 37°C for 1-2 h in buffer containing 500 units of collagenase type IV (Gibco Cat. No. 17104-019) after two consecutive washes with 1% Triton X-100, 0.1M Tris-HCl pH 7.4 buffer and one wash with collagenase buffer (1mM CaCl2, 1% Triton X-100, 0.1M Tris pH 7.4). Following three washes in washing solution, and one hour incubation at room temperature with blocking solution (1% BSA, 0.5% Triton X-100, 1x PBS), the digested animals were stained overnight at 4°C with anti-ALFA primary antibody coupled with the AZDye568 fluorophore (FluoTag-X2 anti-ALFA AZDye568, NanoTag Biotechnologies Cat. No: N1502-AF568-L; dilution 1:100) and diluted in blocking solution. Stained animals were washed thrice with washing solution and then mounted on 5% Noble agar pads in VECTASHIELD antifade mounting medium (Vector Laboratories Cat. No. H-1000) containing 4’,6-diamidino-2-phenylindole (DAPI, 1 mg/ml, Molecular Probes Cat. No. 15451244). Images were acquired on a Leica TCS SP8 microscope and assembled on Fiji version 2.16.0/1.54p (Image J)^77^.

### Alignment of sulfatases and *in silico* signal peptide predictions

For *P. pacificus* and human sulfatases, sequences were accessed from the UniProt platform^78^ and aligned using ESPript 3.0^79^. Sequences of *P. exspectatus* and *P. arcanus* EUD-1 orthologs were retrieved from BLASTp searches on the *Pristionchus* research community public server (http://www.pristionchus.org). The resultant sequences were aligned with *P. pacificus* EUD-1 on Clustal Omega and visualised on Jalview (version 2.11.5.1)^80,81^.

*In silico* signal peptide predictions were carried out on the SignalP 6.0 server^45,46^. The resultant predicted structure and likelihood values were used for data interpretation.

## Data availability

All nematode strains used in this study are available from the corresponding author Ralf J. Sommer. The genome assembly for the *P. pacificus* strain RSC017 and the newly generated whole genome sequencing data of natural isolates have been deposited at the European Nucleotide Archive under the project accession PRJEB108083. All microscopy images are available upon request. Source data are provided with this paper.

## Acknowledgments

We would like to thank relevant La Réunion Island authorities for specimen collection permits and Dr. Matthias Herrmann for the photo of *Amneidus godefroyi*. Dr. Herrmann, Christian Weiler, and former Sommer lab members involved in La Réunion projects are also thanked for help with beetle and nematode collections. We thank Heike Haussmann for freezing of *P. pacificus* materials and Radhika Sharma for discussions. The Weichenrieder lab (Max Planck Institute for Biology, Tübingen) is thanked for providing the Tn5 protein used for tagmentation. This work was supported by the Max Planck Society. P.T. is part of the International Max Planck Research School (IMPRS) ‘From Molecules to Organisms’.

## Author contributions

Conceptualisation: P.T. and R.S.J. Methodology: P.T., H.W., C.R. and C.I. Investigation: P.T., H.W. R.L., T.L., C.R. and C.I. Visualisation: P.T and C.I. Funding acquisition: R.S.J. Project administration: R.S.J. Supervision: R.S.J. Writing—original draft: R.S.J. and C.I. Writing—review and editing: P.T., C.R., C.I. and R.S.J.

---

**Suppl. Fig. 1.**
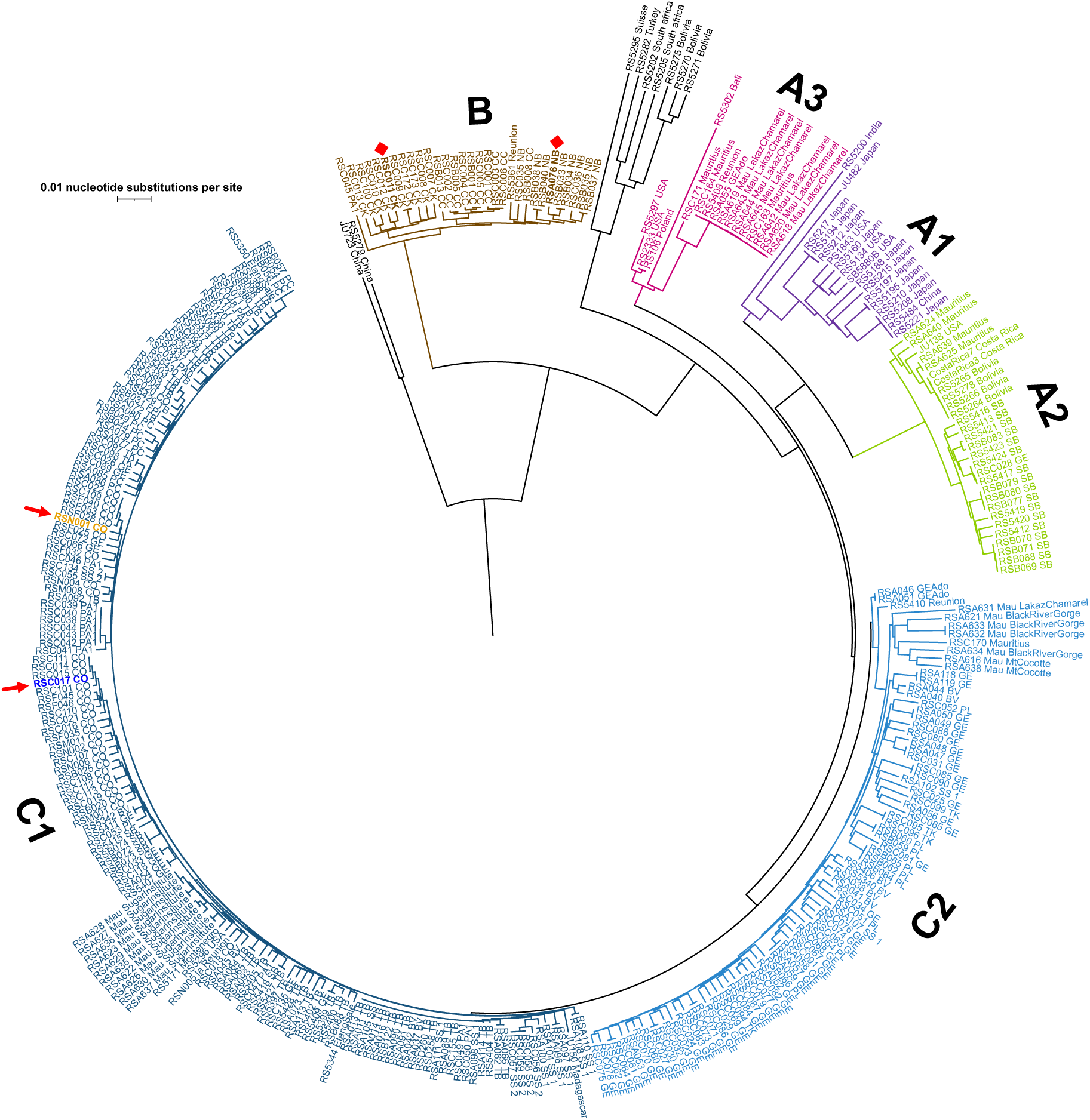
RSC017 and RSN001 strains are closely related. Both *Adoretus*-derived strains (indicated by red arrows) belong to the C1 clade of *P. pacificus* worldwide genotypes. *P pacificus* strains isolated from *Amneidus godefroyi* beetles and used in an earlier study^1^ (indicated by red diamonds) belong to a separate clade: clade B. Scale bar: 0.01 nucleotide substitutions per site. *P. pacificus* clades follow an earlier nomenclature in Rödelsperger et al^2^.

**Suppl. Fig. 2.**
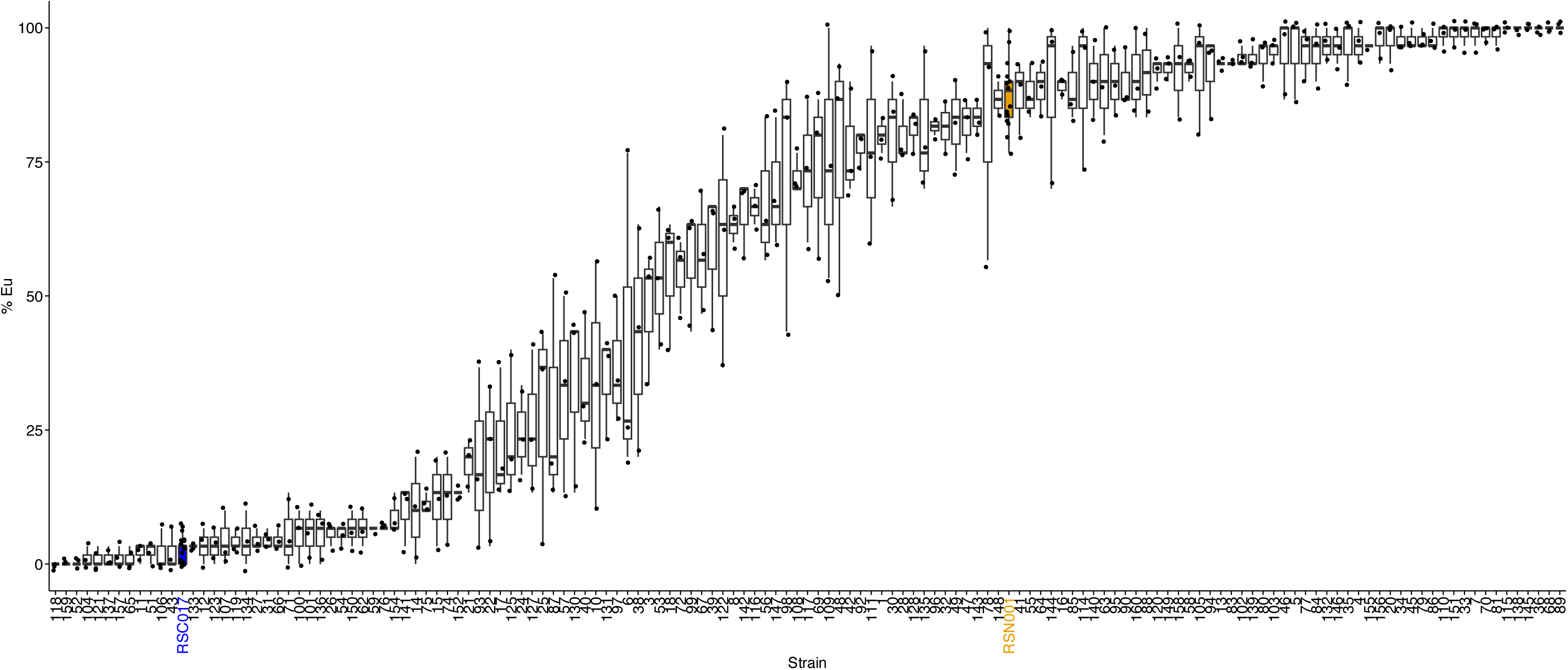
Mouth-form phenotype of RILs and the parental strains RSC017 (St-biased) and RSN001 (Eu-biased). Graph depicts the percentage of Eu animals in the different RILs, RSC017 (in blue) and RSN001 (in orange). Boxplots show the median (centre line), interquartile range (IQR; box), and whiskers extending to 1.5×IQR of mouth-form ratio. Source data are provided as a Source Data file.

**Suppl. Fig. 3.**
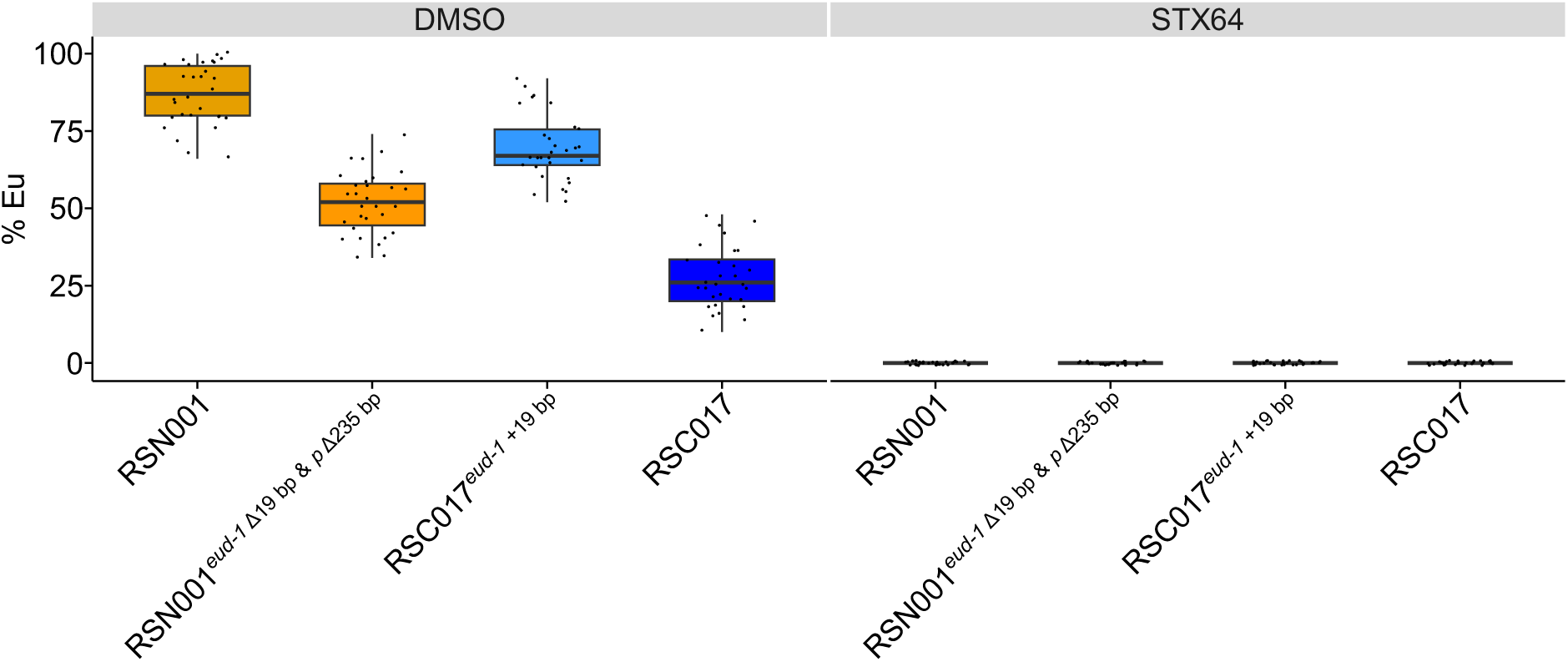
Eu morph in Δ19 bp nonsense variants in exon 1 of *eud-1* relies on EUD-1/sulfatase activity. Quantification of Eu animals in wild type and CRISPR-engineered *eud-1* variants in the absence and presence of the steroid sulfatase inhibitor STX64 (*n* = 30 replicates). Boxplots show the median (centre line), interquartile range (IQR; box), and whiskers extending to 1.5×IQR of mouth-form ratio. Source data are provided as a Source Data file.

**Suppl. Fig. 4.**
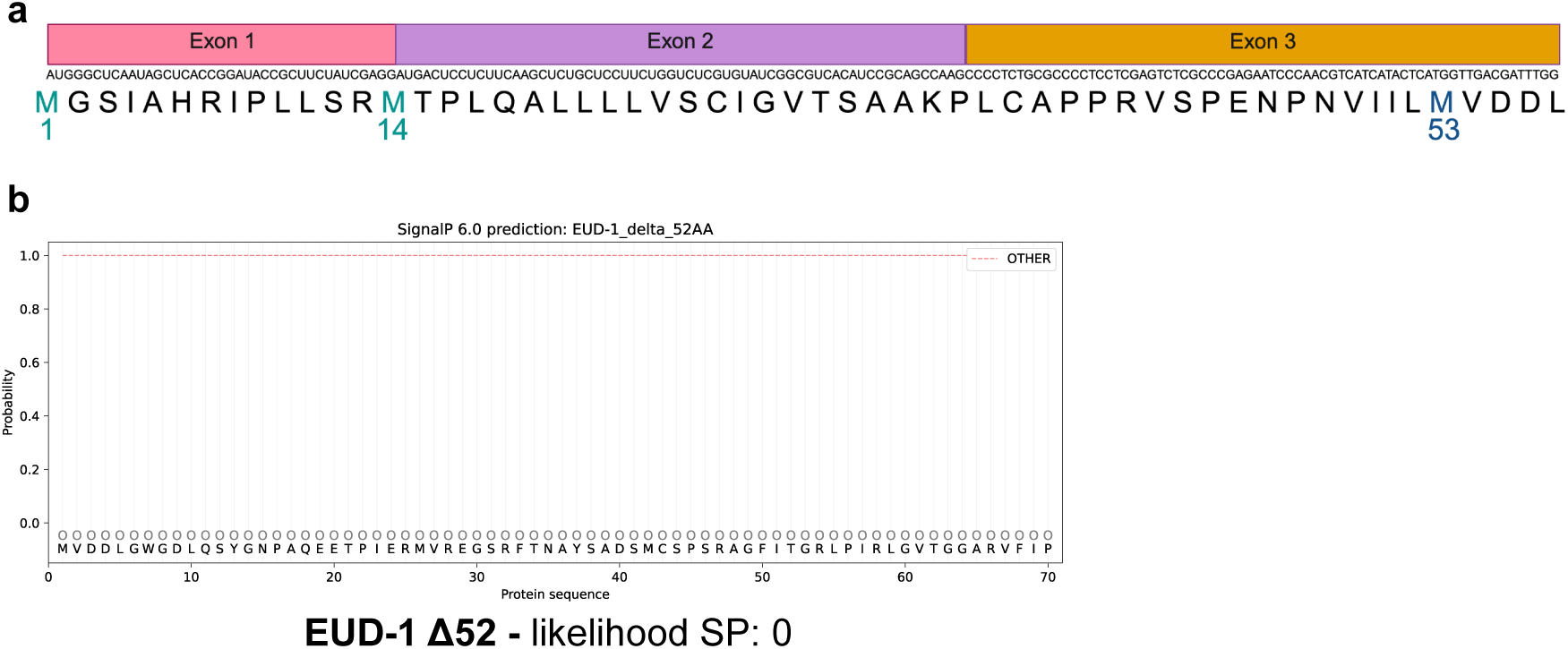
*eud-1* exons 1 and 2 encode a signal peptide. **a**, Schematic representation of *eud-1* exons 1-3 nucleotide and amino acid sequences. Three in-frame Met are present in the first three exons: M1, M14 and M53. **b,** Prediction of signal peptide of a EUD-1 protein lacking the first 52 amino acids using SignalP 6.0. Likelihood value for the prediction is denoted below the graph.

**Suppl. Fig. 5.**
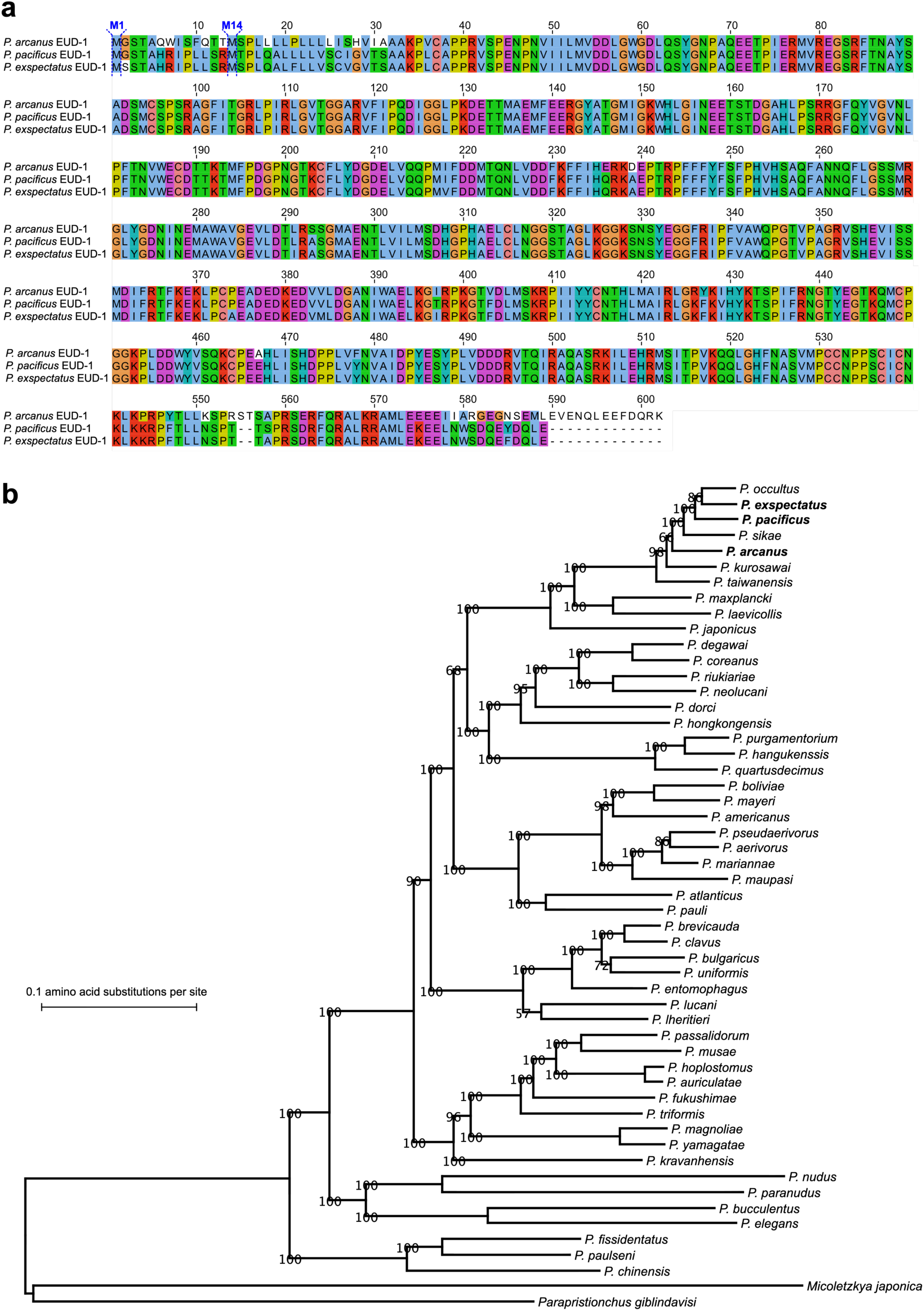
EUD-1 N-terminal proteoforms are present in other *Pristionchus* species. **a**, Alignment of EUD-1 protein sequences across three *Pristionchus* species, *P. pacificus*, *P. arcanus* and *P. exspectatus* highlighting the conservation of M1 and M14 (in blue). **b,** Phylogenetic relationships among *Pristionchus* congeners. *P. pacificus*, *P. arcanus* and *P. exspectatus* are highlighted in bold. Phylogenetic reconstruction adapted from Theam et al^3^. Scale bar: 0.1 amino acid substitutions per site.

